# Sp140L Is a Novel Herpesvirus Restriction Factor

**DOI:** 10.1101/2024.12.13.628399

**Authors:** Jana M. Cable, Wiyada Wongwiwat, Jenna C. Grabowski, Robert E. White, Micah A. Luftig

## Abstract

Herpesviruses, including Epstein-Barr Virus (EBV) – a human oncogenic viruses and essential trigger of multiple sclerosis, must bypass host DNA sensing mechanisms to establish lifelong, latent infection. Therefore, herpesviruses encode viral proteins to disrupt key host factors involved in DNA sensing and viral restriction. The first viral latency protein expressed, EBNA-LP, is essential for transformation of naïve B cells and establishment of viral gene expression, yet its role in evading host defenses remains unclear. Using single-cell RNA sequencing of EBNA-LP-Knockout (LPKO)- infected B cells, we reveal an antiviral response landscape implicating the ‘speckled proteins’ as key cellular restriction factors countered by EBNA-LP. Specifically, loss of *SP100* or the primate-specific *SP140L* reverses the restriction of LPKO, suppresses a subset of canonically interferon-stimulated genes, and restores transcription of essential latent viral genes and cellular proliferation. Notably, we also identify Sp140L as a restriction target of the herpesvirus saimiri ORF3 protein, implying a role for Sp140L in immunity to other diverse DNA viruses. This study reveals Sp140L as a restriction factor that we propose links sensing and transcriptional suppression of viral DNA to an IFN-independent innate immune response, likely relevant to all nuclear DNA viruses.

**Significance Statement:** Herpesviruses, including the oncogenic Epstein-Barr virus (EBV), are restricted by DNA sensing during initial infection and therefore encode viral proteins to antagonize key restriction factors. We found that the EBV latency protein EBNA-LP, disrupts the ‘speckled proteins’ Sp100 and Sp140L - an evolutionarily recent protein with unknown function, which we find promotes an anti-viral state that suppresses cellular proliferation, characterized by high induction of cellular anti-viral genes and suppressed transcription of essential viral latency genes. Sp140L also restricts the herpesvirus saimiri, which we find antagonizes Sp140L through the viral protein ORF3. Our study therefore identifies Sp140L as a novel restriction factor of diverse herpesviruses, and likely all DNA viruses, during a critical stage of initial viral infection.

## Introduction

Herpesviruses are double stranded DNA viruses that cause lifelong infections due to their ability to establish viral latency and avoid immunodetection (1). Clinically relevant herpesviruses include herpes simplex virus (HSV), human cytomegalovirus (HCMV) and Kaposi’s sarcoma associated herpesvirus (KSHV), as well as Epstein-Barr virus (EBV), which is associated with numerous lymphomas and carcinomas, and is implicated in several autoimmune disorders – particularly multiple sclerosis (2-4). The B cell biology of EBV infection and associated diseases can be modeled *in vitro,* as infection of primary B cells leads to cellular transformation and the generation of immortalized lymphoblastoid cell lines (LCLs). Given the ability of herpesviruses to establish lifelong latency, initial infection is a crucial stage where cellular sensing and restriction can prevent the virus from successfully establishing latency. Yet, the mechanisms of intrinsic restriction of herpesviruses have not been fully elucidated.

One key mediator of defense against DNA viruses are PML nuclear bodies (PML-NBs), membrane-less nuclear compartments involved in the intrinsic DNA sensing and epigenetic repression of viral chromatin through loading repressive histones and epigenetic modifications on the incoming naked viral DNA genomes (5, 6). Components of PML-NBs can also play a role in the innate antiviral response through inducing expression of interferon-stimulated genes (ISGs) (7-12). Core PML-NB proteins implicated in restricting DNA viruses include PML, DAXX, ATRX, and Sp100. PML serves as a scaffold to mediate the interaction of other core proteins, while DAXX and ATRX together load the repressive histone variant H3.3 onto viral genomes during initial infection - preventing the accumulation of active histone marks on the incoming viral genome and suppressing the transcription of viral genes (13-17). HIRA, a separate histone chaperone complex recruited to PML-NBs upon herpesvirus infection (17) and IFN treatment (8, 17), also contributes to H3.3 deposition on viral chromatin in HSV infection (17, 18). In contrast, H3.3 loading by HIRA at ISG gene loci activates gene expression (7, 8). The speckled protein Sp100, can restrict the replication and viral gene expression of the DNA viruses HSV (19, 20), HCMV(21), Human Papilloma Virus (22), and adenovirus (23) when these viruses lack effective countermeasures or when their countermeasures are species-mismatched. Sp100 can bind the heterochromatin protein HP1α (24) and can localize to the promoters of ISGs upon IFN stimulation (10). However, the mechanisms by which Sp100 restricts DNA virus infection is still unclear.

Beyond *SP100*, the speckled protein genetic locus includes additional family members *SP110*, *SP140*, and – exclusively in primates – *SP140L* (25), whose roles in viral infection are largely unknown. The speckled family genes all encode four key domains: i) an N-terminal caspase recruitment (CARD) domain (previously called heterogeneously staining region – HSR) involved in multimerization, ii) the DNA-binding (SAND) “Sp100, AIRE, NucP41/P75 and DEAF” domain, iii) a methyl-histone-binding plant homeodomain (PHD) domain, and iv) a C-terminal acetyl-lysine-binding bromodomain (BRD) (25). Genetic mutations in *SP110* and *SP140* are associated with autoimmune disorders and increased susceptibility to intracellular bacterial infections (25). And while Sp110 and Sp140 repress the Type I IFN response in mice (26, 27), they appear to promote Type I IFN responses in humans (28). The *SP140L* gene arose as a likely meiotic crossover event between the *SP100* and *SP140* genes, resulting in a gene combining the 5’ CARD domain-encoding region of *SP100* and the remaining protein-encoding domains of the *SP140* gene (29). The function of Sp140L, however, remains unexplored.

To overcome PML-NB restriction, successful herpesviruses encode proteins that antagonize various PML-NB components of their hosts. These include viral tegument proteins belonging to the viral FGARAT homolog family, such as pp71 in HCMV (30, 31) and BNRF1 in EBV (32) which target DAXX and ATRX; ORF75 in KSHV which antagonizes DAXX, ATRX, PML, and Sp100 (33), and ORF3 of Herpesvirus Saimiri (HVS) – a KSHV-like rhadinovirus of squirrel monkeys – which degrades Sp100 (34). Some immediate-early viral proteins also antagonize PML-NBs, including ICP0 in HSV - a viral E3 ubiquitin ligase which targets PML, Sp100, DAXX, and ATRX (35, 36); IE1 in HCMV targeting PML and Sp100 (21, 37); and EBNA-LP in EBV, which transiently displaces Sp100 from PML-NBs during early infection (38, 39). While the molecular mechanism by which these viral proteins antagonize PML-NB components may be distinct, the functional consequences are similar as evidenced by the ability of ICP0 or various combinations of BNRF1 or pp71 with IE1 or EBNA-LP to complement an ICP0-null HSV and pp71-deficient HCMV (40-42). The role of Sp100 in EBV infection has not been directly investigated, with studies largely limited to using transient transfection rather than primary infection systems (38, 39), and focusing on the shortest Sp100 isoform – Sp100A – which does not contain the SAND, PHD, or Bromo domains, and promotes viral gene transcription, whereas the longer Sp100 isoforms restrict DNA viruses (23, 43).

EBNA-LP is critical for the transformation of naïve B cells by EBV infection(44), but its function in early infection is not well defined compared to other PML-NB-disrupting herpesvirus proteins mentioned above. The architecture of EBNA-LP is unique in that it is composed of N-terminal, identical, 66 amino acid repeats (W domains) and a C-terminal 45 amino acid Y domain. While the structure of EBNA-LP is unknown, it is predicted to be highly disordered(45). Based on transient transfection assays, EBNA-LP was thought to primarily function as a co-activator of another early EBV protein, EBNA2 to enhance expression of the essential viral gene Latent Membrane Protein 1 (LMP1) and some cellular genes (46-50). The mechanism by which EBNA-LP co-activates EBNA2 has been variously attributed to displacement of the transcriptionally repressive HDAC4 and NCoR proteins from viral and cellular promoters (51, 52). However, B cells infected with an EBNA-LP Knockout EBV (LPKO) exhibit reduced expression of not only LMP1, but also other viral genes that are not EBNA2-dependent, while EBNA2-induced host gene expression is not reduced, or even increased, in the absence of EBNA-LP during infection(44) suggesting that EBNA-LP’s relationship to EBNA2 is more complex than a simple ‘coactivator’. Additionally, our recent work identified highly conserved, hydrophobic leucine-rich motifs in both the W and Y domains of EBNA-LP that associate with the DNA-looping factor YY1 (53). Thus, EBNA-LP likely has EBNA2-independent roles that are critical for establishing latency in, and transformation of, naïve B cells.

Given the essential role of immediate-early herpesvirus proteins in overcoming intrinsic viral restriction, and particularly the gap in knowledge as to how and why EBNA-LP is necessary for EBV infection and transformation of naïve B cells, we used single cell RNA-sequencing (scRNAseq) to compare the trajectories of wild type (WT) and LPKO virus infected B cells during the first days of infection. This assay revealed key facets of the restriction of EBV in the absence of EBNA-LP. We then defined the cellular factors and mechanisms responsible for the restriction of the LPKO virus, and finally the breadth of how these factors are restrictive across human DNA viruses.

## Results

### EBNA-LP Dictates the Fate Trajectories of EBV-Infected B Cells

To assess the role of EBNA-LP in EBV-driven B cell fate trajectories, we performed single cell RNA sequencing (scRNAseq) of primary human B cells prior to infection (Day 0), and at 2, 5, and 8 days following WT and LPKO infection as depicted in **Fig. 1A**. B cell purity (**Fig. S1A**) and infection efficiency (**Fig. S1B**) were confirmed by expression of CD19 and CD23 respectively at time of collection. Single cell RNA-seq revealed that infected samples contained more mRNA (**Fig. S1C**) and more unique mRNA features (**Fig. S1D**) per cell than uninfected (Day 0) B cells. Dimensional reduction integrating all seven samples generated a single UMAP for further analysis (**Fig. 1B**). Uninfected cells largely cluster separately from infected cells (**Fig. 1C**) and 2 days post-infection LPKO- and WT-infected clusters are nearly identical to each other (**Fig. S1D and S1E**). Differences in cellular states between WT- and LPKO-infected B cells emerge by 5 days (**Fig. S1F and S1G**) and are accentuated 8 days post-infection (**Fig. S1H and S1I**).

**Fig. 1.**
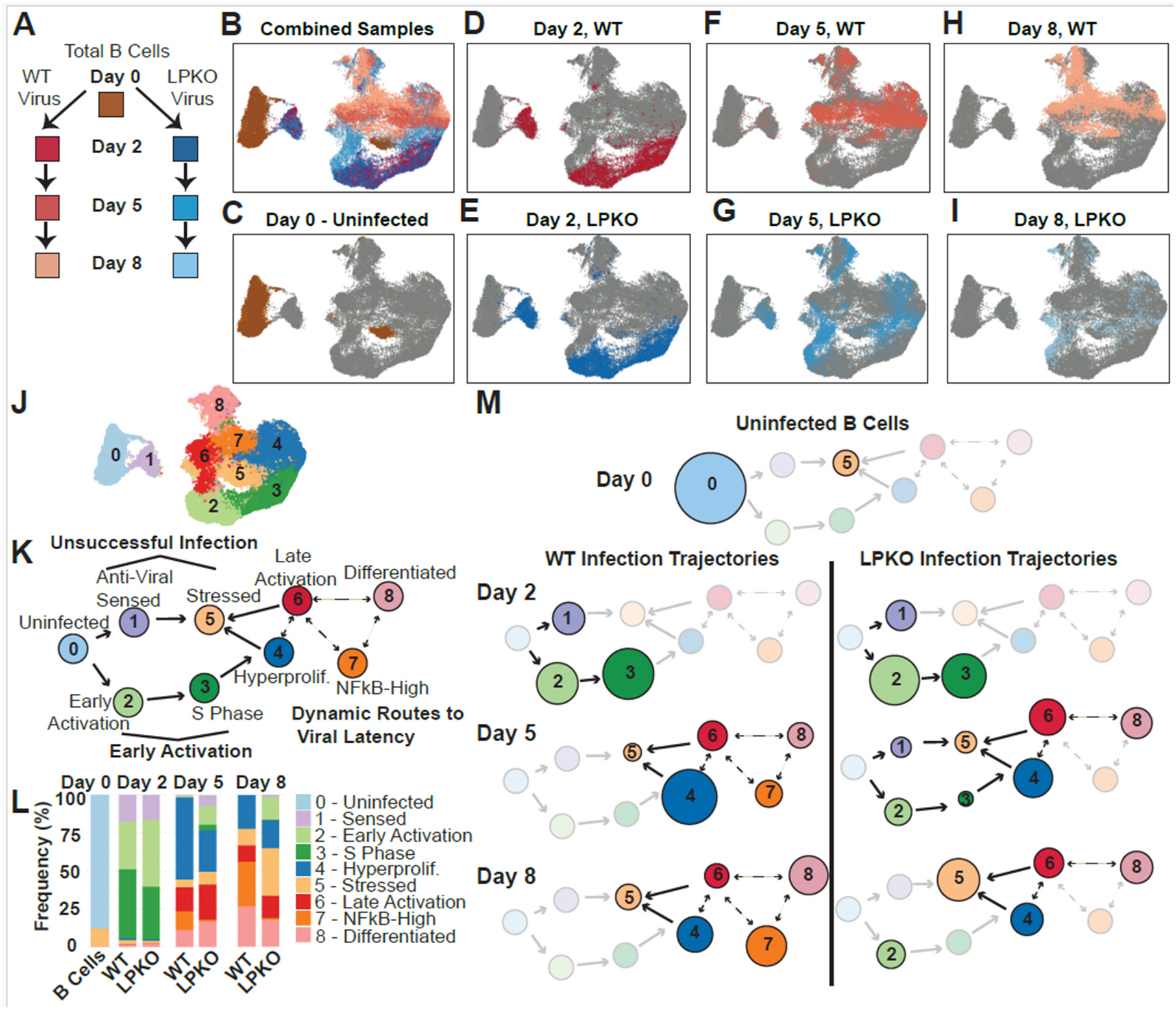
LPKO-infected B cells have altered cellular trajectories compared to WT EBV. (**A**) Schematic of collected samples. (**B**) Uniform Manifold Approximation and Project for Dimension Reduction (UMAP) of the combined seven samples. Colors correspond to samples in (**A**). (**C**) UMAP with uninfected B cells highlighted. (**D**) Cells 2 days post-infection with WT. (**E**) Cells 2 days post-infection with LPKO. (**F**) Cells 5 days post-infection with WT. (**G**) Cells 5 days post-infection LPKO. (**H**) Cells 8 days post-infection WT. (**I**) Cells 8 days post-infection LPKO. (**J**) UMAP with cells grouped into 9 unique clusters based on differential gene expression. (**K**) Model of the identified clusters in the cellular trajectories of EBV infected B cells. (**L**) Frequency of each cluster within each sample. (**M**) Model of the trajectories of WT- and LPKO-infected B cells at each time point based on frequencies in (**L**). Circle area is proportional to frequency of cluster in each sample. Transparent circles represent clusters which are absent.

Our prior work defined a set of distinct B cell fate trajectories following EBV infection that largely mimic the states of B cell activation and germinal center maturation(54). In comparing WT- and LPKO-infected cells, we have identified 9 distinct cell states that largely recapitulated those characterized previously (**Fig. 1J and Table S1**). We used the top cluster-defining genes in our previous study to identify the corresponding cell states in this dataset (**Fig. S2A-S2I**). Cell states were further distinguished on the basis of viral gene expression (**Fig. S2J-S2U**) and cell cycle phase (**Fig. S2V-S2X**). Cluster 0 (Uninfected – c0) contained uninfected naïve and memory B cells (**Fig. 1K**). Two days post-infection, prior to cellular proliferation, which begins around 3 days post-infection (55), a portion of cells exist in a gene expression state associated with antiviral sensing (Sensed – c1) (**Fig. 1K**). Cells that avoid this initial barrier transition towards an activated state, resembling pre-germinal center B cells (Early Activation – c2), and towards a state of DNA replication enriched for cells in S phase (S Phase – c3) (**Fig. 1K**). By 5 days post-infection, cells advanced to a state of hyperproliferation, with markers of dark zone germinal center B cells (Hyperproliferating – c4) (**Fig. 1K**). Likely as a result of the cellular stress associated with EBV-induced hyperproliferation (55, 56), by this time point some cells also arrest in a state of cellular stress (Stressed – c5) (**Fig. 1K**). Cells also transition towards a second, but distinct, activated B cell state (Late Activation – c6) (**Fig. 1K**). Compared to the Early Activation state (c2), this Late Activation state (c6) is more highly enriched in naïve B cells based on IgD expression (**Fig. S2X**), has lower expression of Myc and E2F targets (**Fig. S2Y and S2Z**) likely a consequence of reduced expression of the viral transactivator EBNA2 in c6 compared to c2 (57) (**Fig. S2R**), and is likely dependent upon cells having undergone proliferation based on its appearance only after day 2 (**Fig. 1K**). EBV-infected cells also enter a NFκB-High state, likely driven by the viral protein LMP1, whose expression is highest in this population (**Fig. S2T**), that is characterised by markers of light zone germinal center B cells (NFκB High - c7) (**Fig. 1K**). The final population is a differentiated plasmablast-like state (Differentiated - c8) (**Fig. 1K**).

While this model held true for WT-infected cells, LPKO-infected cells had altered trajectories. First, unlike WT-infected cells, LPKO-infected cells remain in the early (pre-proliferation) states of activation (c2 and c3) beyond 2 days post infection (**Fig. 1L**). Second, LPKO-infected cells were also more likely than WT-infected cells to enter states of antiviral sensing (c1) or cellular stress (c5) after day 2 (**Fig. 1L**). Finally, the LPKO-infected cells largely failed to establish the NFκB-High state (c7) (**Fig. 1L**), consistent with the previously reported low expression of LMP1 in LPKO infections (44) (**Fig. S2U**). Surprisingly, the LPKO virus efficiently established the plasmablast state (Differentiated – c8) (**Fig. 1L**), despite the absence of the NFκB-High state which is critical in the emergence of plasmablasts from the germinal center (58). Interpreted as trajectories in **Fig. 1M**, LPKO-infected cells persist in early infection states of activation (c2 and c3) and cellular stress prior to proliferation (c1) longer than WT. LPKO-infected cells that do hyperproliferate, have a higher propensity to arrest with signs of cellular stress (c5) and to enter a germinal center- or light zone-independent route towards the plasmablast state (c8), similar to germinal center-independent B cell maturation (59).

To validate our transcriptome data, we first identified RNA changes in cell surface markers defining the key cell populations. RNA levels of CCR6 (**Fig. 2A**) and CD23 (**Fig. 2B**) in the scRNA-seq data distinguish many cell states, similar to previous studies(54). Early Activation states (c2 and c3) are characterized by CCR6^hi/^CD23^lo^ expression, while the Late Activation state (c6) is CCR6^hi/^CD23^hi^. The NFκB-High state (c7) is CCR6^lo^/CD23^hi^ and is further resolved by high expression of ICAM1 (**Fig. 2C**), a proxy marker for LMP1 expression (60). The remaining states are CCR6^lo^/CD23^lo^ with the Stressed (c5) distinguishable from the Differentiated state (c8) by low expression of CD38 (**Fig. 2D**) and reduced cellular proliferation. Therefore, these markers along with the proliferation tracker CellTrace should allow quantitation of the main states that differ between LPKO and WT infections: persistence in states of activation (c2, c3, c6), reduced transition to the NFκB-high state (c7), and increased cellular arrest prior to hyperproliferation (c5) as outlined in **Fig. 2E**.

**Fig. 2.**
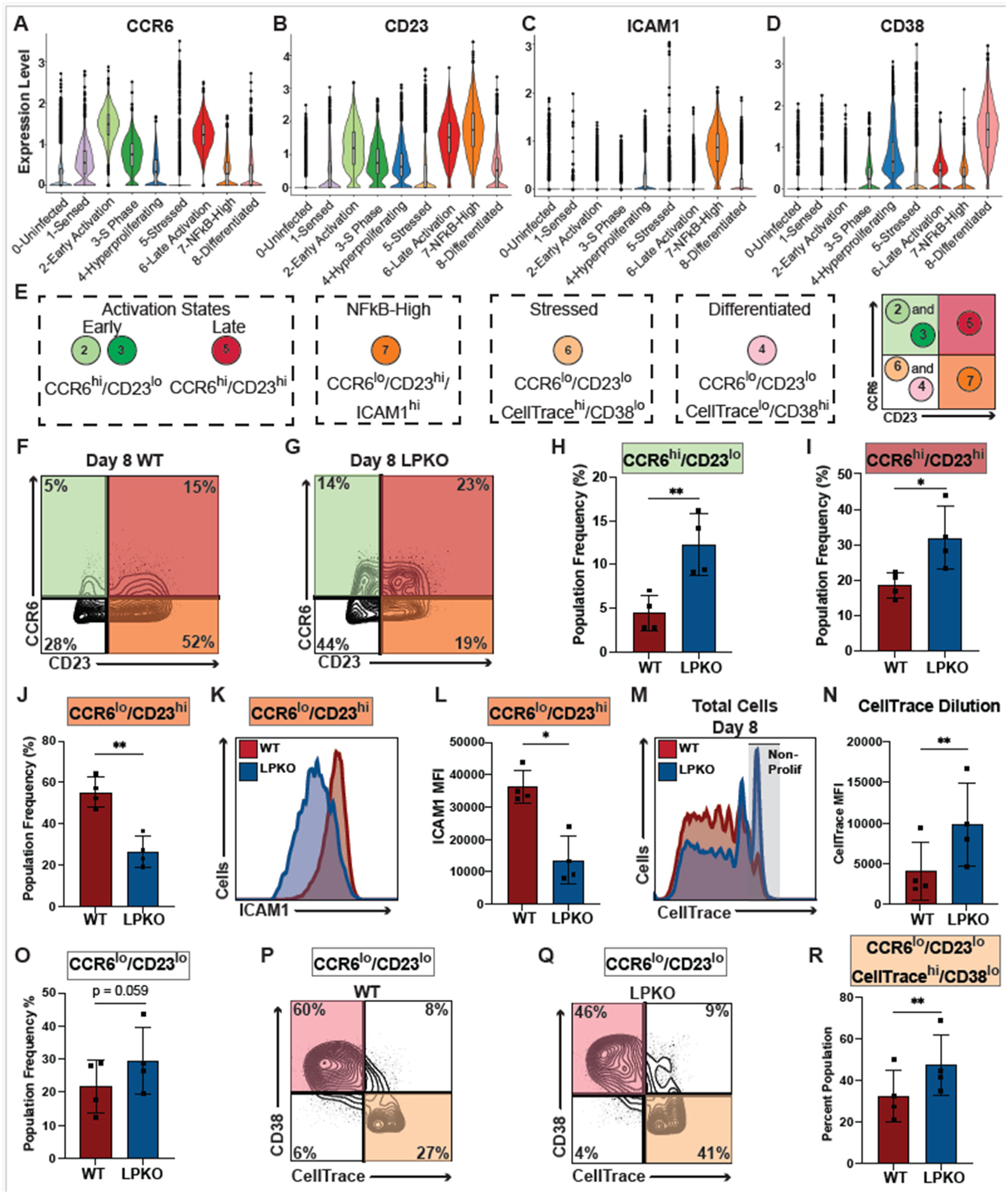
Altered cell populations between WT- and LPKO-infected B cells identified in scRNAseq data are validated by flow cytometry 8 days post-infection. Expression levels of CCR6 (**A**), CD23 (**B**), ICAM1 (**C**), and CD38 (**D**) in scRNA-seq clusters. (**E**) Legend of flow panel marks for the key states identified. Flow staining for CCR6 and CD23 8 days post-infection in representative Donor 2 for WT (**F**) and LPKO (**G**) infected B cells. Quadrant color corresponds to cell states that can be distinguished as depicted in (**E**). Percent of WT- and LPKO-infected cells 8 days post-infection in the CCR6^hi^/CD23^lo^ (Early Activation) states (**H**), the CCR6^hi^/CD23^hi^ (Late Activation) state (**I**), and the CCR6^lo^/CD23^hi^ (NFκB-High) states (**J**). (**K**) Histogram comparing expression of ICAM1 protein (proxy for LMP1) between WT and LPKO infected cells in CCR6^lo^/CD23^hi^ population. (**L**) Quantified Median Fluorescence Intensity (MFI) of ICAM1 signal in CCR6^lo^/CD23^hi^ population in WT and LPKO infections (n = 4). (**M**) Histogram of CellTrace stain, diluted during each cell division, 8 days post infection in WT and LPKO infected cells in representative Donor 1. Grey area indicates cells that have not proliferated. (**N**) MFI of CellTrace (n=4). Lower MFI indicates more cell proliferation. (**O**) Percent of WT and LPKO infected cells 8 days post-infection in the CCR6^lo^/CD23^lo^ states (n=4). Separation of CCR6^lo^/CD23^lo^ populations in WT (**P**) and LPKO (**Q**) infection by CD38 and CellTrace to distinguish Differentiated cells from Stressed cells. (**R**) Percent of CCR6^lo^/CD23^lo^ cells that are also CD38^lo^ and CellTrace^hi^ (indicating low proliferation) representing the stressed cell cluster. Significance determined with ratio paired t test. N = 4 biological replicates. *Indicates p value less than or equal to 0.05, ** indicates p value less than or equal to 0.01. Histograms are scaled as percent of maximum count (modal).

Using Flow cytometry to analyze protein expression of identified markers in a new set of infected samples, we classified cells by CCR6 and CD23 levels (**Fig. 2F, 2G, S3A, and S3B**) and confirmed the prolonged presence of LPKO-infected cells in states of early activation (c2 and c3) (CCR6^hi^/CD23^lo^) (**Fig. 2H**) and late activation (c6) (CCR6^hi^/CD23^hi^) (**Fig. 2I**) compared to WT infection. Second, we confirmed that LPKO-infected cells have reduced transition to NFκB-high state (c7), as LPKO-infected cells have reduced CCR6^lo^/CD23^lo^ populations (**Fig. 2J)** and the cells that do enter this state express significantly less ICAM1 **(Fig. 2K, 2L, and S3C)** compared to WT. Furthermore, compared to WT, LPKO-infected cells are less proliferative (**Fig. 2M, 2N, and S3D).** And, upon further separating CCR6^lo^/CD23^lo^ cells based on CD38 expression and proliferation, we confirmed that compared to WT, LPKO-infected cells are more frequently in states of cellular stress (c5) (CCR6^lo^/CD23^lo^/CellTrace^hi^/CD38^lo^) (**Fig. 2O-2R, S3E, and S3F**). Altogether, these data support a model in which EBNA-LP is important in preventing the arrest, stress response, and immune activation after EBV infection.

### LPKO-Infected Cells Fail to Suppress the Antiviral Response and Fail to Induce Cellular Metabolic Remodeling

We next sought to characterize the molecular mechanisms underlying the cell fate restrictions following LPKO infection. We first pooled all time points for each WT and LPKO scRNAseq dataset into a “pseudo-bulk” dataset and performed gene set enrichment analysis (GSEA) to identify differentially regulated pathways between WT and LPKO infection. In WT-infected cells, we observed significant enrichment of transcripts involved in NFκB signaling (**Fig. 3A**) in agreement with the failure of LPKO-infected cells to generate an NFκB-high state (**Fig. 1L**). We also observed an increase in expression of genes controlling cellular metabolism in WT-relative to LPKO-infected cells (**Fig. 3A**). In contrast, LPKO-infected cells were enriched for transcripts of interferon-stimulated genes (ISGs) and antiviral responses (**Fig. 3B**).

**Fig. 3.**
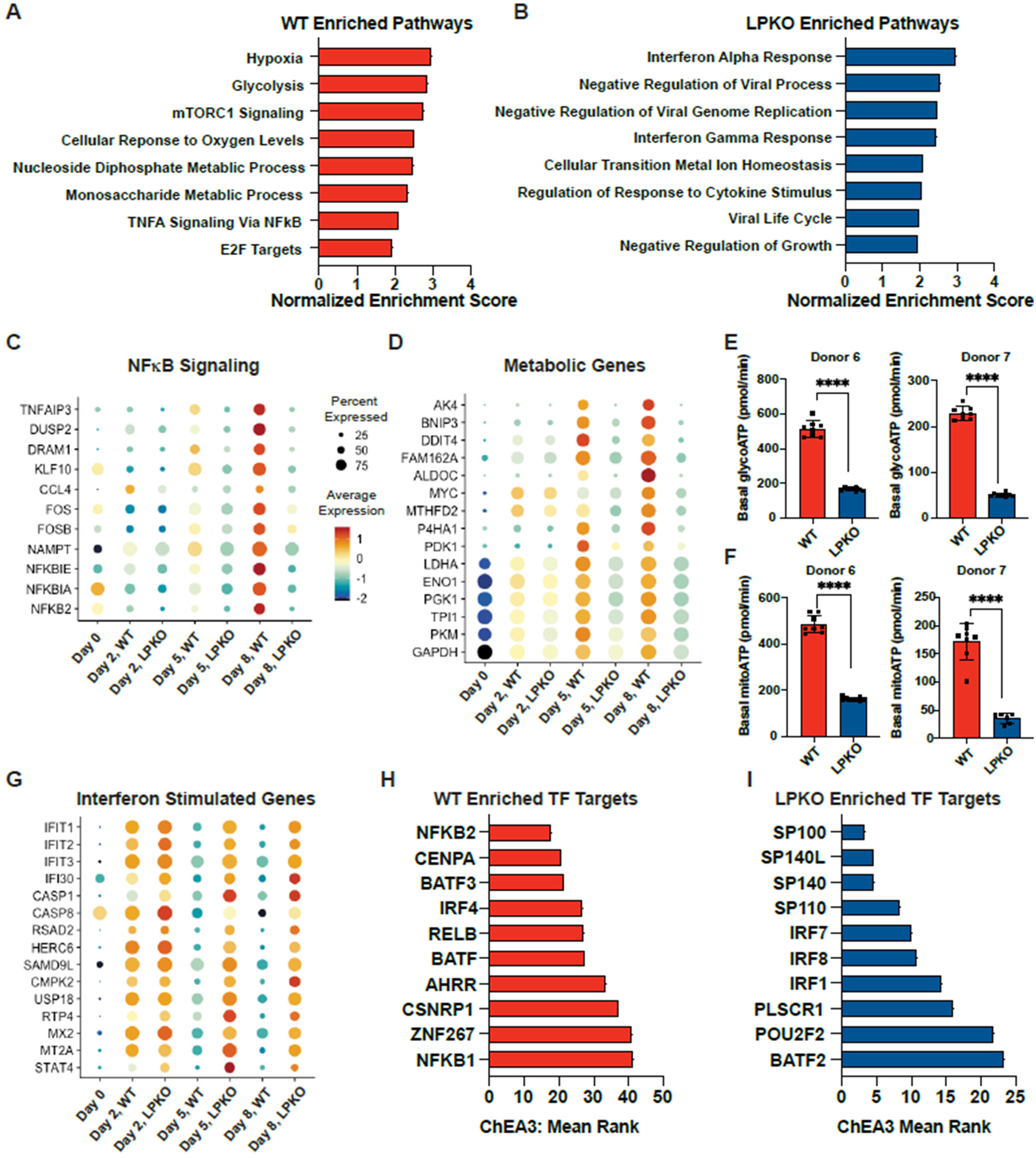
EBNA-LP promotes expression of pro-proliferative genes, while reducing expression of anti-viral genes. Pathways identified as enriched in WT- compared to LPKO- (**A**) or LPKO- compared to WT- (**B**) infected scRNAseq samples by GSEA Hallmark gene sets or Gene Ontology datasets. Representative top-ranking pathways with false discover rates (FDR q-values) less than 0.05. Higher normalized enrichment score indicates stronger enrichment. Dot plot of representative NFκB signaling-related genes (**C**) and metabolism genes (**D**) identified in GSEA pathways enriched in WT infected cells for each sample in the scRNAseq time course. (**E**) Basal rate of ATP derived from glycolysis (glycoATP) in WT- and LPKO-infected naïve B cells 4 days post-infection in two donors. (**F**) Basal rate of ATP derived from oxidative phosphorylation (mitoATP) in WT- and LPKO-infected naïve B cells 4 days post-infection in two donors. (**G**) Dot plot of representative Interferon Stimulated Genes enriched in LPKO-infected cells for each sample in the time course. (**H**) Top ten predicted transcription factors identified by ChEA3 associated with expression of the genes enriched in WT-infected cells in scRNAseq samples compared to LPKO or (**I**) LPKO-infected cells compared to WT. Lower mean rank indicates a stronger correlation.

We next used time-resolved pseudo-bulk analysis to better define the dynamic nature of EBNA-LP-dependent gene regulation. NFκB targets were modestly decreased 2 days post-infection in both WT and LPKO-infected cells, but then diverged at 5, and more so at 8, days post infection (**Fig. 3C**). Cellular metabolic gene expression diverged by day 5, following the first EBV-induced cell division (**Fig. 3D**). While loss of NFκB signaling is concurrent with failure to efficiently express LMP1 (44) (**Fig. S2U**), failure to upregulate cellular metabolism in the absence of EBNA-LP was previously unexplored. Therefore, we further examined cellular metabolic activity during early infection and confirmed naïve B cells infected with LPKO virus had reductions in both basal glycolysis and oxidative phosphorylation compared to WT infected cells at 4 days post-infection (**Fig. 3E and 3F**).

Transcripts downregulated upon expression of EBNA-LP (i.e. induced in LPKO-infected cells) were primarily ISGs. Intriguingly, though, interferon gene mRNAs were not elevated in LPKO-infected cells (**Fig. S4A**), indicating ISG induction may occur through other mechanisms. These ISG transcripts are widely induced by two days post-infection in both WT- and LPKO-infected cells (**Fig. 3G**). In the presence of EBNA-LP, WT-infected cells effectively down-regulated expression of these ISGs, whereas in LPKO-infected cells, levels of these ISG transcripts were sustained or increased (**Fig. 3G**). These trends in differentially expressed pathways observed in the scRNAseq data were confirmed by bulk RNAseq of WT- and LPKO-infected cells in additional donors at 8 days post-infection (**Fig. S4B-S4F**).

Because the ISG induction appeared to be independent of IFN induction, we sought to identify which cellular factors could be mediating the observed gene expression differences in WT- and LPKO-infected cells. We used ChEA3 (61) (ChIP-X Enrichment Analysis Version 3) to identify transcription factors whose target genes were enriched in the transcriptome of WT- and LPKO- infected cells. Genes upregulated in WT-infected cells were associated with factors including NFκB subunits; CENPA – a centromere protein and transcriptional regulator that forms a complex with YY1 (62) which we recently identified as an EBNA-LP-interacting protein (53); and IRF4, which can suppress ISG transcription (63), induce expression of Myc (63, 64), and function in cooperation with BATF and BATF3 that are also suggested mediators of these expression changes by the analysis (65, 66) (**Fig. 3H**). Genes upregulated in the absence of EBNA-LP were instead associated with all four human speckled proteins (Sp100, Sp110, Sp140, and Sp140L) (**Fig. 3I**).

### Sp100 and Sp140L, but Not Other PML-NB Proteins, are Required to Restrict EBV Transformation in the Absence of EBNA-LP

To examine whether the ChEA3-identified speckled proteins or other PML-NB proteins specifically restrict EBV infection in the absence of EBNA-LP, we assessed the effect of knocking out these target genes on rescuing the inability of LPKO virus to transform naïve B cells. Naïve B cells isolated from adult peripheral blood (**Fig. S5A**) were directly transfected with Cas9-guide RNA ribonucleoprotein (Cas9/RNP) complexes targeting genes encoding the main components of PML bodies, including the speckled proteins, along with CD46, a non-essential cell surface protein used as a proxy for knockout of the target gene (67) (**Fig. 4A**). Transfected cells were then infected with either the LPKO or WT virus, and outgrowth of edited, CD46-negative cells was assessed by flow cytometry weekly post-infection, for 4 weeks (**Fig. 4A**). As expected, control LPKO-infected cells in which only *CD46* was targeted failed to sustain proliferation to generate LCLs, whereas WT EBV still transformed the naïve B cells into LCLs (**Fig. 4B-4D**). While knockout of most of the PML-NB-associated proteins that were targeted – including *PML* – had no significant impact on the outgrowth of LPKO-infected cells, which decreased in number over time like the *CD46*-only knockout, mutation of either *SP100* or – even more emphatically – *SP140L* rescued outgrowth (**Fig. 4B and 4C**). Indeed, knockout of *SP100* or *SP140L* was sufficient to consistently generate LPKO LCLs by 28 days post-infection (**Fig. 4D**) and target genes were efficiently knocked out in these LCLs as validated by genomic DNA sequencing (**Fig. S5B**) and protein expression where applicable (**Fig. S5C**). Additional LPKO LCLs were generated after a longer time in culture with inconsistent target knockout (**Fig. S5D**), likely arising from contaminating memory B cells. Targeting the same factors in WT infected cells had no impact on outgrowth (**Fig. S5E and S5F**), confirming Sp100 and Sp140L restrict EBV infection only in the absence of EBNA-LP.

**Fig. 4.**
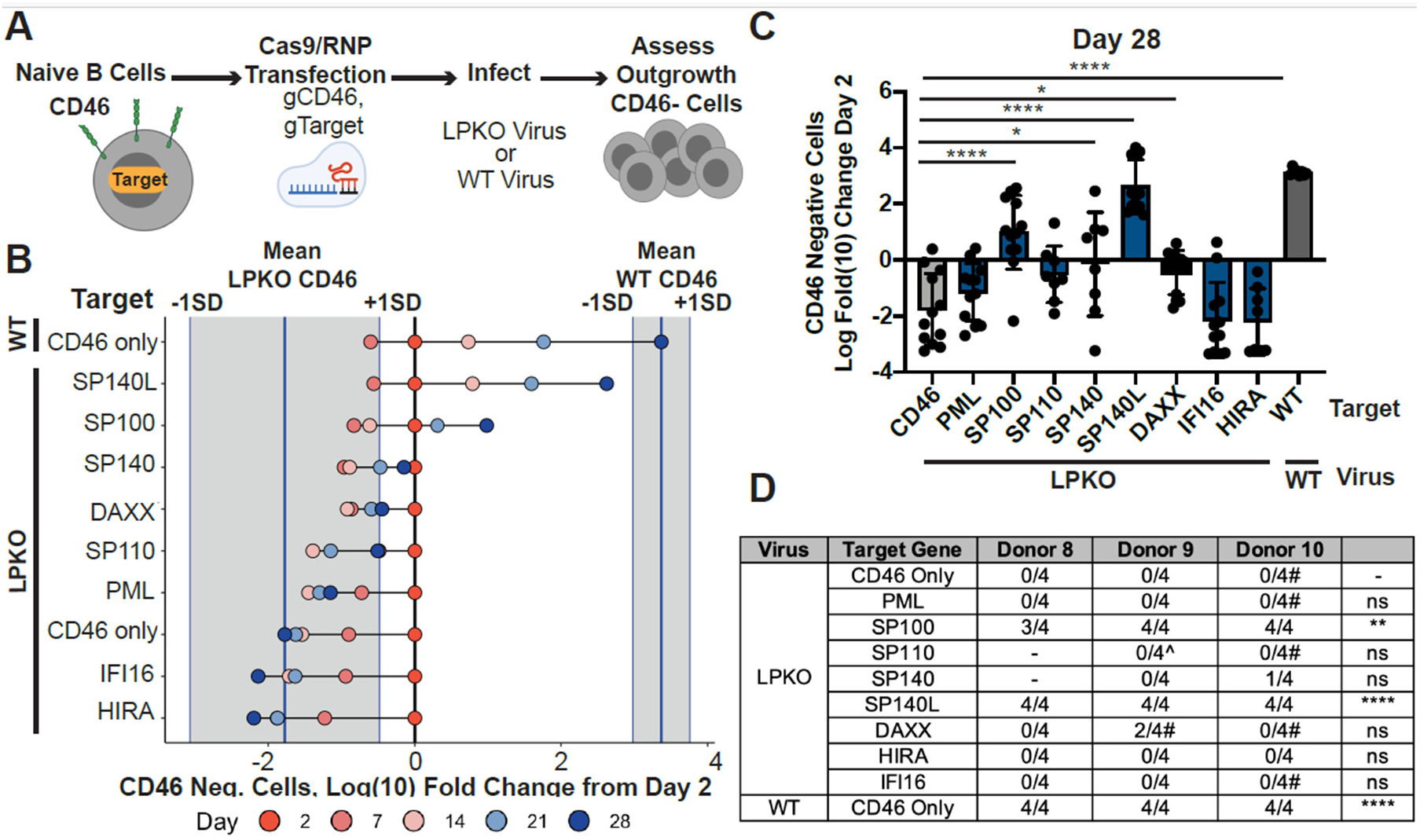
Knockout of *SP140L* or *SP100* rescues transformation of LPKO-infected naïve B cells. (**A**) Schematic of Cas9/RNP screening approach. (**B**) Log(10) Fold Change from Day 2 of the total number of CD46 negative cells for each condition 2, 7-, 14-, 21-, and 28-days post-infection. Dark turquoise line represents the mean number of CD46 negative cells in CD46 only control WT- or LPKO-infected cells 28 days post infection. SD = standard deviation. (**C**) Total number of CD46 negative cells 28 days post infection for each condition plotted as log fold change from 2 days post infection. P values calculated by one-way ANOVA with multiple comparisons to compare to LPKO infected cells transfected with guide targeting CD46 only as control. *Indicates p-values <0.05, ** indicates p-values <0.01, **** indicates p value < 0.0001. (**D**) Total number of LPKO LCLs in each donor for each target out of 4 replicates per donor. ^#^Indicates conditions in which at least one LCL was generated after an additional three weeks in culture with variable knockout efficiency. ^Indicates conditions in which at least one LCL was generated in the same time frame at WT virus, but target gene was not knocked out. P values calculated using Fischer’s exact test to compare outcomes to LPKO control condition ** indicates p-values <0.01, **** indicates p value < 0.0001.

### Structural characterization of EBNA-LP/Sp140L and Sp100 interactions define a critical interface important for B cell transformation

To better understand the nature of the putative interaction between EBNA-LP and Sp140L and Sp100, we used Alphafold3 to predict the structure of a complex between these proteins (68). First, we found that despite EBNA-LP being predicted to have high intrinsic disorder (45), AlphaFold3 consistently predicted that EBNA-LP forms an extended beta sheet, with loops connecting helices derived from the repeated W domains and a loop extending the unique 45 amino acid Y domain containing a short alpha helix (**Fig. 5A and 5B**). The interaction sites with Sp140L and Sp100 are consistently predicted to involve this Y domain alpha helix engaging a CARD domain derived helical bundle of Sp140L and Sp100 (**Fig. 5C and 5D**), the latter being consistent with prior biochemical findings (38). Specifically, the hydrophobic interface of these interactions relies on a conserved leucine rich motif (LRM) within EBNA-LP (53) (**Fig. 5E and 5F**). Indeed, modification of the leucines in this helix to alanines was sufficient to disrupt the association between EBNA-LP and Sp100 by co-immunoprecipitation (**Fig. 5G**).

**Fig. 5.**
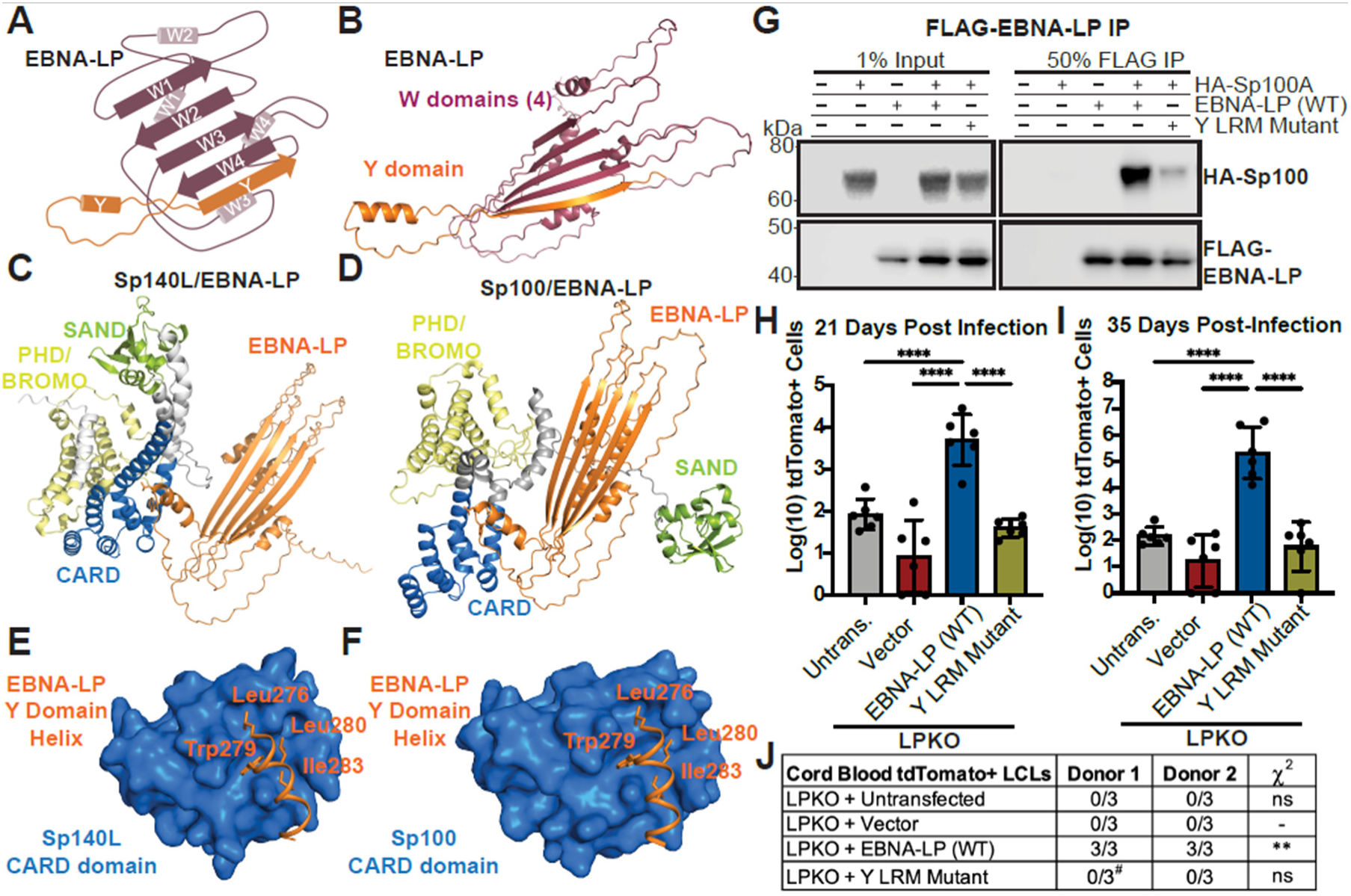
EBNA-LP is predicted to engage Sp140L and Sp100 through an alpha-helix containing a leucine-rich motif in the C-terminal Y domain. (**A**) Schematic of EBNA-LP (encoding 4 repeated W domains) predicted structure W (magenta) and Y domain (orange) contributions to beta strands and alpha helices based on **B**. (**B**) Alphafold3-predicted model of EBNA-LP three-dimensional structure. Predicted model of EBNA-LP (orange) in complex with (**C**) Sp140L and (**D**) in complex with Sp100. Speckled protein domains are highlighted: CARD domain (blue), SAND domain (green), PHD/BROMO domain (yellow). Zoomed in view of EBNA-LP Y domain helix (spanning amino acids 276 through 287) encoding the leucine-rich motif in contact with (**E**) Sp140L and (**F**) Sp100 CARD domains. EBNA-LP residues with side-chains predicted to contact CARD domains are indicated. **G**. FLAG co-immunoprecipitation of FLAG-tagged EBNA-LP wild type or mutant with leucines in Y domain alpha-helix modified to alanine (Y LRM Mutant, L276,280,281A) with HA-Sp100 Isoform A performed in 293T cells. EBNA-LP constructs encode 4 W domains. 1% of input material and 25% of co-immunoprecipitation was used for imaging. (**H**)Total tdTomato positive cells in each condition of cord blood B cells infected with LPKO and trans-complemented with indicated DNA construct 21 days post-infection and (**I**) 35 days post-infection (performed in two donors, with 3 replicates each). Untransfected cells indicates background signal threshold. P values calculated by ordinary one-way ANOVA with Dunnett’s multiple comparisons test. **** Indicates p value < 0.0001. (**J**) Number of tdTomato positive LCLs generated per condition. ^#^ Indicates a single replicate in which cells were transformed but were not tdTomato positive. P values were calculated using Fisher’s exact test to compare transformation outcomes to LPKO-infected cells trans-complemented with empty vector. ** Indicates p value < 0.01.

To further examine the role of the Y domain LRM at the speckled protein interface in transformation of naïve B cells, we used a trans-complementation assay in cord blood B cells infected with LPKO virus as previously described (53). Prior to infection, cells were transfected with an episomal plasmid encoding tdTomato and either wild type or an EBNA-LP mutant in which leucines were mutated to alanine. We first confirmed expression of the mutant protein in LPKO-transformed total B cells, since EBNA-LP is not required for the transformation of the memory B cell component, and demonstrated that both wild type and mutant EBNA-LP are able to be expressed (**Fig. S6A**) and had the same sub-cellular localization profile as EBNA-LP expressed from WT virus (**Fig. S6B**). EBNA-LP with a mutated Y domain LRM failed to rescue transformation of LPKO-infected cord blood B cells compared to wild-type EBNA-LP (**Fig. 5H – 5J**) – indicating that the Y domain LRM of EBNA-LP involved in Sp100 and likely Sp140L interaction is essential during EBV infection.

### *SP100* or *SP140L* Knockout Rescues Cellular and Viral Gene Expression Critical for Naïve B Cell Outgrowth

We then sought to investigate the mechanism by which Sp100 and Sp140L restrict EBV infection in the absence of EBNA-LP. We used bulk RNAseq to examine the effect of *SP100* and *SP140L* knockout on viral and cellular gene expression five days after LPKO infection of naïve B cells (**Fig. S7A**). Knockout of *SP100* or especially *SP140L* reduced induction of IFN-related genes in LPKO-infected cells (**Fig. 6A-6D**). Knockout of either *SP100* or *SP140L* also rescued the expression of pathways important for proliferation including MYC targets, E2F targets, and particularly upon *SP140L* knockout, metabolic pathways (**Fig. 6A-6C and 6E**). NFκB signaling was also rescued by *SP100* and *SP140L* knockout in LPKO infected cells (**Fig. 6A-6C and 6F**), with knockout of *SP140L* again exhibiting in a more complete rescue of gene expression towards wild-type levels – in agreement with the transformation screen (**Fig. 4B**). Since LMP1 and other viral genes are inefficiently activated during LPKO infection (44), we examined viral latency gene expression, and found that *LMP1* and *LMP2* RNA levels in LPKO infected cells were somewhat elevated upon loss of *SP100* and largely recovered in *SP140L* knockouts (**Fig. 6G**). This was further confirmed at a functional level by flow cytometry using ICAM1 as a proxy for LMP1, as knockout of *SP100* or – to a greater extent – *SP140L* rescued surface ICAM1 protein to levels comparable to WT infected cells (**Fig. 6H and S7B**).

**Fig. 6.**
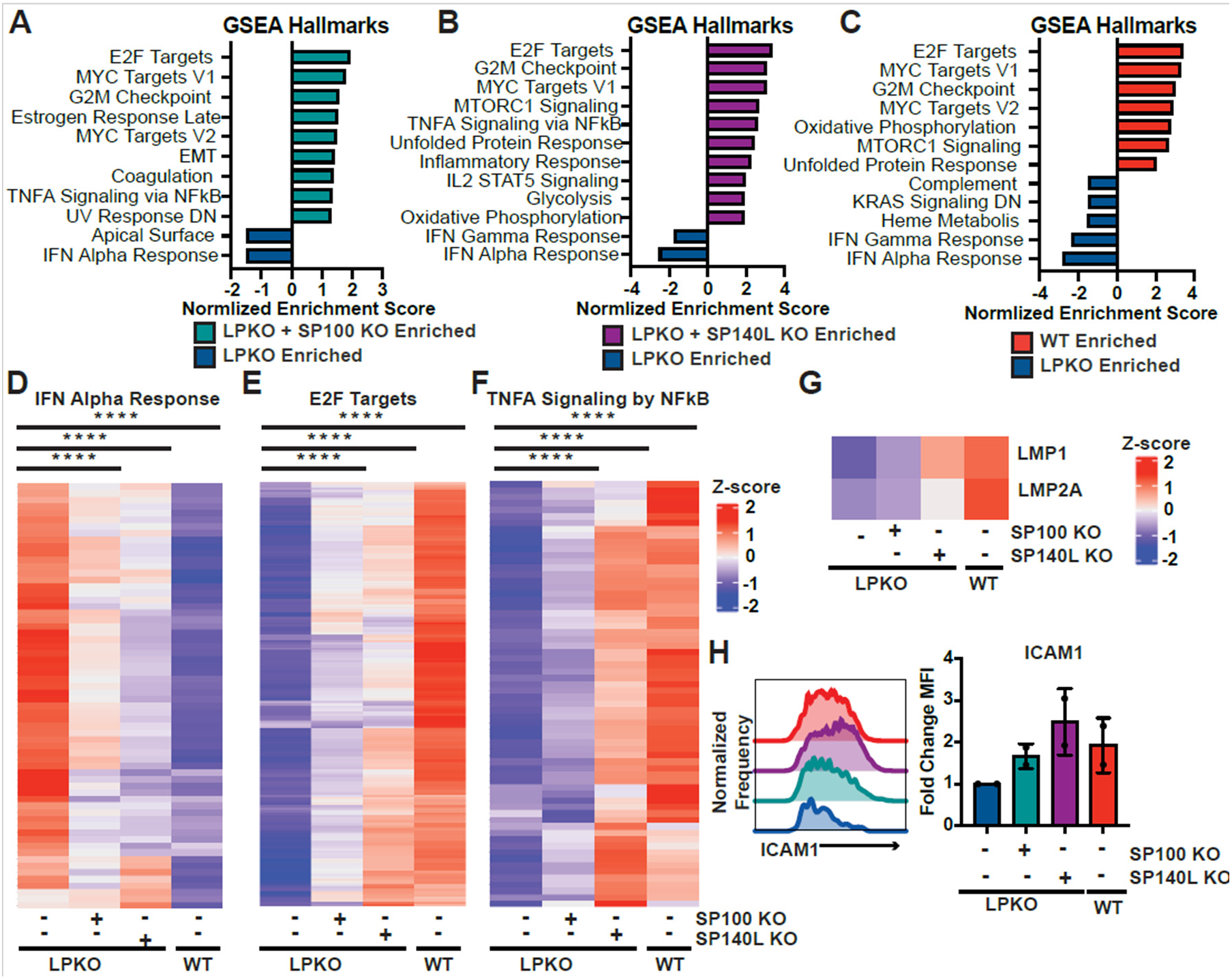
*SP100* and *SP140L* Knockout in LPKO-infected cells promotes transcription of pathways required for outgrowth and inhibits the anti-viral response. GSEA Hallmark pathway analysis of bulk RNA-sequencing samples 5 days post-infection comparing LPKO infected cells with (**A**) *SP100* knockout, (**B**) *SP140L* knockout or WT virus infection (**C**) to in comparison control LPKO infection (n = 2). GSEA terms with nominal p value less than 0.05 are shown. Higher absolute value of the normalized enrichment score indicates stronger enrichment. Heatmap of genes related to the significantly enriched Hallmark pathways (**D**) “Interferon (IFN) Alpha Response,” (**E**) “E2F Targets”, and (**F**) “TNFA Signaling by NFκB” for each condition. Average z-score of two samples is plotted. P values determined by One-way Repeated Measures ANOVA with multiple comparisons to LPKO control. **** indicates p value less than 0.0001. (**G**). Average Z-score for mRNA encoding LMP1 and LMP2A for each sample. (**H**). Histogram of flow cytometry signal for LMP1 proxy gene ICAM1 in each sample.

### Sp140L Also Restricts ORF3-Deficient Herpesvirus Saimiri (HVS)

As Sp140L has not previously been studied as a viral restriction factor, we sought to determine whether Sp140L can restrict herpesviruses other than EBV, and in cell types other than B cells, by studying Sp140L in HVS infection. As the HVS tegument protein ORF3 targets Sp100 for degradation(34), we generated two independent WT HVS and ORF3-KO HVS viruses (**Fig. S8A and 8B**) carrying a CMV promoter-driven luciferase expression cassette. We confirmed infection of human fibroblast HFF cells with WT HVS, but not ORF3-KO HVS, resulted loss of Sp100 protein (**Fig. S8C**). We then compared virus transcription after initial infection in two human fibroblast cell lines (HFF and MRC5 cells) carrying knockout of either *SP100* or *SP140L* alongside the proxy gene *CD46* (**Fig. 7A**). Consistent with a previous report with an ORF3-KO of a C-strain HVS (34), knockout of *SP100* rescued viral transcription after ORF3-KO HVS infection (**Fig. 7B and 7C**). Consistent with our EBV data, knockout of *SP140L* was also sufficient to rescue luciferase expression from the ORF3-KO virus in both cell types (**Fig. 7B and 7C**). Loss of *SP100* or *SP140L* in the context of WT HVS infection had little or no effect on viral transcription (**Fig. S8D-S8F**) suggesting that ORF3 antagonizes both Sp100 and Sp140L function during WT HVS infection. These findings suggest Sp140L is a restriction factor of diverse gammaherpesviruses in primates, that is at least as important as Sp100.

**Fig. 7.**
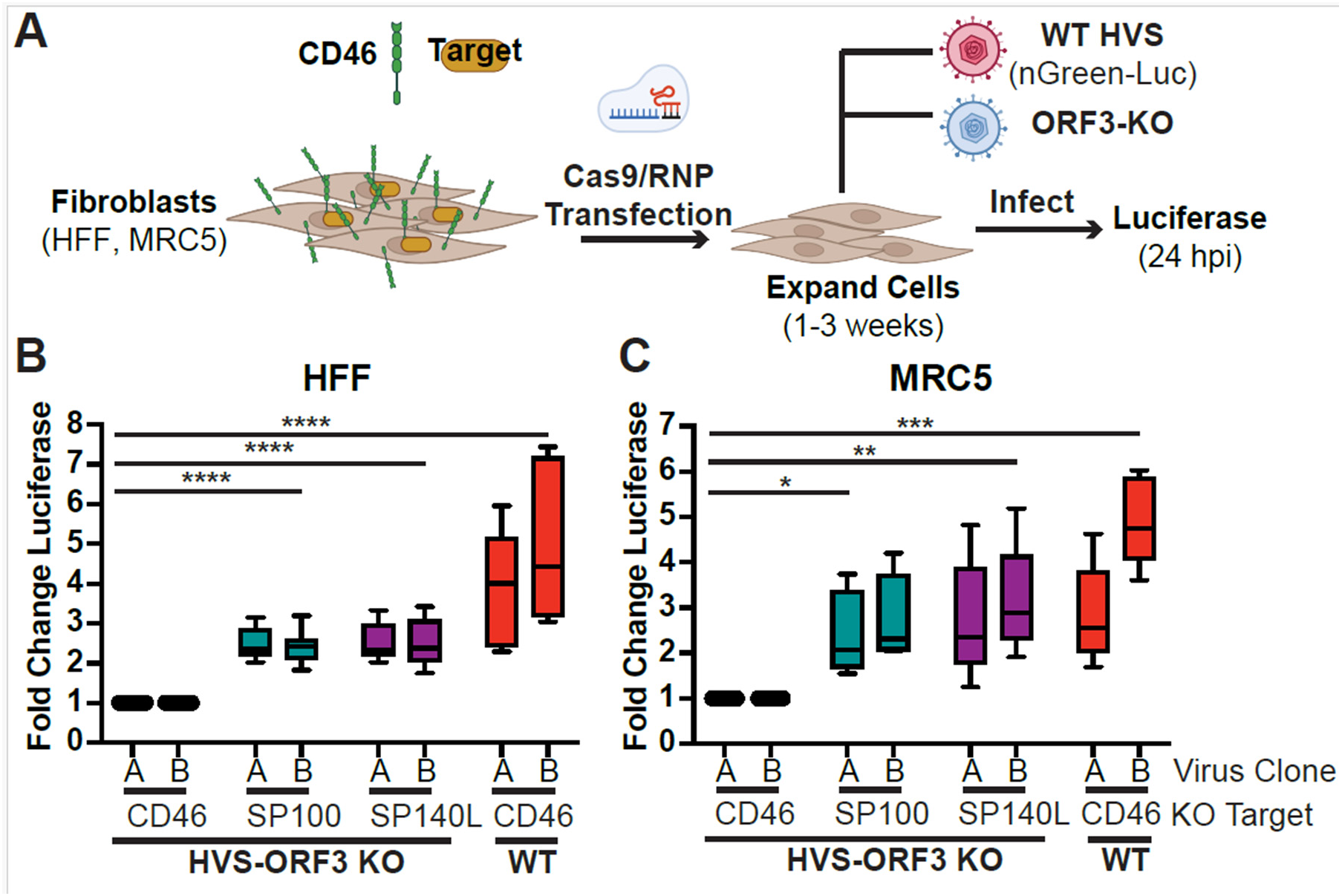
Sp100 and Sp140L restrict Herpesvirus Saimiri (HVS) in the absence of ORF3. (**A**). Schematic of experiment. Fold change in luciferase expression HFF cells (n=7) (**B**) or MRC5 cells (n=5) (**C**) infected with HVS-ORF3 Knockout or WT virus encoding luciferase with knockout of control *CD46*, *SP100*, or *SP140L*. Two virus preparation (A and B) were used. P values calculated by Dunnett’s multiple comparisons test using an ordinary two-way ANOVA. * Indicates p-values <0.05, ** indicates p-values <0.01, **** indicates p value < 0.0001.

## Discussion

Herpesviruses infect nearly all animal species and result in lifelong latent and persistent infections (69). As DNA viruses, herpesviruses must overcome cellular barriers to establish infection, including the detection and chromatinization of their foreign DNA genomes by intrinsic restriction factors such as those in PML-NBs, including Sp100. Here, we have identified Sp140L, an Sp100-homolog, as a novel restriction factor of herpesvirus infection that is counteracted by the EBV latency protein EBNA-LP and the HVS tegument protein ORF3. Our data are consistent with a mechanistic model whereby both Sp100 and Sp140L restrict EBV infection by preventing the expression of the essential viral latency genes that are expressed after EBNA-LP and EBNA2, and by altering cellular gene expression - both inducing an ISG response and reducing B cell metabolic capacity. While the core antiviral PML-NB proteins have been thought to include PML, Sp100, DAXX, and ATRX, our work adds the evolutionarily recent Sp140L to the list of key cellular intrinsic restriction factors of DNA viruses in primates.

Leveraging a targeted CRISPR screen, we identified Sp100 and Sp140L as key restriction factors that prevent EBV transformation of naïve B cells in the absence of EBNA-LP and viral transcription of HVS in the absence of ORF3. Since knocking out other PML-NB components did not rescue LPKO EBV, these other components of PML-NBs must either be fully targeted by other EBV proteins, exhibit functional redundancy, or are not required to block EBV-mediated transformation. Based on studies of HSV and CMV mutants that lack abilities to inhibit PML-NB functions (13, 30, 32, 70, 71), we can conclude that PML-NBs have two parallel repressive pathways in humans: one driven by DAXX/ATRX and another by Sp100 and – we now propose – Sp140L. In the LPKO virus, its DAXX-counteracting tegument protein BNRF1 is still intact, suggesting that factors in our screen that failed to rescue naïve B cell transformation may be part of the DAXX-associated pathway. While DAXX/ATRX load repressive histones (e.g. H3.3) onto the incoming viral genome (13), Sp100 and Sp140L may be responsible for the addition of repressive histone marks such as H3K9me3 and/or H3K27me3. For example, during latent KSHV infection of endothelial cells, loss of Sp100, but not DAXX nor PML, correlates with H3K27me3 deposition on the viral genome (72), although Sp100’s ability to bind heterochromatin protein 1 (HP1) and high-mobility group protein B1 (HMGB2) (73) suggests Sp100 may also be linked to H3K9me3 (74).

While PML-NB components are known to restrict DNA virus gene expression, we discovered that Sp100 and Sp140L are also important in suppressing cellular proliferation. Possible mechanisms of restricting proliferation include the failure to express the LMPs during LPKO infection, however, LMP1 is not essential for the initial cell divisions after EBV infection (75). An alternative mechanism explaining these results is that these factors integrate intrinsic DNA sensing with an anti-proliferative ISG response. Recent evidence suggests Sp100 can localize to ISG promoters (10) suggesting Sp100 – and perhaps by extension Sp140L – induce ISGs through direct transcriptional regulation or chromatin modification at ISG promoters - as has been observed with other PML-NB components including HIRA (7, 8), ATRX(9), and PML (11) in mutant HSV- and HCMV-infected cells or upon IFN stimulation. We also observed a restoration of expression of proliferation-linked genes upon knockout of *SP100* or – more convincingly – *SP140L*, suggesting that detection of foreign DNA or upregulation of ISG proteins may repress pro-proliferative genes. For example, RSAD2, an ISG highly expressed during LPKO compared to WT EBV infection, influences cellular metabolism and mitochondrial function (76, 77). IRF proteins can also target metabolic genes (78, 79), and notably several IRFs were also identified as potential mediators of gene changes in LPKO-infected cells by ChEA3 (**Fig. 3G and 3H**). In contrast, whether Sp100 and Sp140L impose cellular arrest by functioning as cell cycle checkpoint proteins, therefore resulting in loss of proliferation-associated genes remains unknown. Still, these findings suggest Sp100 and Sp140L impose an antiviral state through not only regulating key viral genes, but also through repression of cell cycle progression to impose an anti-proliferative state not favorable for viral infection or transformation.

To further investigate the mechanism by which EBNA-LP may disrupt Sp100 and Sp140L, we used AlphaFold3 to generate the first predicted structure of EBNA-LP and found that an alpha helix in the C-terminal Y domain of EBNA-LP is predicted to associate with the CARD domain of Sp100 and Sp140L. Mutagenesis of the leucine-rich motif (LXXXLL) within this alpha helix disrupts EBNA-LP’s interaction with Sp100 and likely Sp140L, and reduces the ability of EBNA-LP to trans-complement the LPKO virus in the transformation of naïve B cells. Our previous work identified this motif as important for EBNA-LP to associate with the cellular transcription factor YY1 (53). This motif is similar in sequence to both the LXXLL motif found in many cellular proteins that regulate transcription (80), and to a similar but more diverse in sequence motif termed “flexiNR” that is encoded in many viral proteins with transcription regulation functions (81). Our findings fit with the notion that many viral proteins, including viral oncoproteins, have evolved short linear motifs, generally less than 10 residues long in which a few key residues define affinity and specificity, in order to mimic and/or hijack cellular protein functions (82, 83). One such example includes the LXCXE motif in the many viral proteins that can promote S Phase transition through interaction with pRb, p107, and p130 (84). Therefore, this LXXXLL motif is an example of short linear motif critical for viral protein function.

This study was strengthened by the use of primary cells rather than cancer cell lines. This is critical as PML-NBs functions can be disrupted in cancer cells, and therefore the use of cancer cell lines for DNA sensing or virus restriction assay may not reflect physiological relevance (8). Still, there are limitations to performing knockouts in primary B cells, as to reduce cell death prior to infection, B cells are infected with EBV only 2 hours post-transfection. Therefore, as EBNA-LP is one of the earliest expressed EBV latency proteins, turnover of targeted protein may be incomplete by the time EBNA-LP would be expressed, even if genomic editing has occurred, and therefore this assay cannot fully complement the LPKO virus. While the HVS assay using primary fibroblasts to generate knockout cell lines prior to infection does not have the same temporal disadvantage, it was challenging to get complete knockout of target genes even in cells with the proxy target, CD46, lost. While overcoming these limitation would likely allow for an even more pronounced rescue of the LPKO and ORF3-KO viruses with knockout of *SP100* and *SP140L*, our assays clearly show that both *SP100* and *SP140L* restrict these diverse herpesviruses. Furthermore, in our assays, knockout of *SP140L* resulted in stronger phenotypic rescue – supporting Sp140L as the predominant speckled protein restricting DNA viruses. However, we cannot exclude the possibility these observations are a result of differences in knockout efficiency between *SP100* and *SP140L*.

The *SP140L* gene was formed by a gene duplication and cross-over event between *SP100* and *SP140* shortly prior to the emergence of primates (29). As a result, the Sp140L N-terminal CARD domain is over 90% identical to that of Sp100, while its C-terminal SAND and PHD/Bromodomains are very like those of Sp140 (29). Sp140L also lacks the exons that in Sp100 encode SUMOylation and HP1 binding motifs but retains the exons that in Sp100 can bind to the high mobility group protein HMGB2. Given the CARD domain is responsible for the dimerization(24, 85) or multimerization(86) of Sp100 and that Sp100 and Sp140L colocalize in cells (29), and that Sp100 can form homotypic interactions with the more distantly related protein Sp110 (87), we anticipate that Sp100 and Sp140L can interact, perhaps interchangeably, through this domain. This is further supported by our findings that knockout of either Sp100 or Sp140L alone was sufficient for rescue of transformation. Whether Sp140L arose as the result of an evolutionary arms race with viral antagonists of Sp100 remains a possibility. As knockout of Sp140 did not enhance transformation of either WT- or LPKO-infected B cells, further analysis of the molecular differences between Sp140L and Sp140 – and the key domains responsible for DNA virus restriction requires further investigation. Additionally, expression differences between Sp100 and Sp140L may contribute to the dependency of EBNA-LP in naïve, but not memory B cell transformation as at the RNA level, naïve B cells more highly express both Sp100 and Sp140L (88).

Based on what is known about Sp100, we can propose the following model for how both Sp100 and Sp140L might sense and suppress incoming viral genomes. The SAND domain of Sp100 is reported to preferentially bind pairs of unmethylated CpG dinucleotides (89). Its PHD/Bromodomain contains bulky residues where other PHD/Bromodomains contain binding pockets for histone modifications (90). CARD domains commonly multimerize through homotypic interactions to activate immune signaling pathways (91). Speckled proteins Sp100, Sp110 (87), and Sp140 (92, 93) also associate on cellular chromatin at gene promoters including Sp100A at ISGs (10), with Sp140 localizing to sites enriched in H3K27me3 to maintain heterochromatin (92, 93). We hypothesize therefore that the SAND domain of Sp100 (and by extension Sp140L) recognize the unmethylated incoming viral DNA as foreign, perhaps in combination with the PHD/Bromodomain recognizing features of newly assembled chromatin. Multimerization of Sp100 and Sp140L proteins along the viral genome can then block the initiation of transcription of viral genes (with only immediate early genes having escaped repression) either directly or by recruiting repressors like HP1 or other chromatin remodeling factors to promote or maintain the viral genome heterochromatin. Meanwhile, viral infection may also influence Sp100 and/or Sp140L localization to cellular gene promoters, perhaps relieving repressive epigenetic marks at the loci of ISGs. Alternatively, multimerization of the CARD domains nucleates a signaling process that activates transcription of ISGs (87), activating cell division checkpoints to suppress proliferation of a cell that has potentially been invaded by a persistent foreign genome.

In conclusion, we identify Sp140L as a restriction factor of herpesviruses, and likely all DNA viruses, targeted by critical viral proteins of primate viruses, but may represent a significant barrier to any virus with a nuclear DNA phase of its lifecycle that transmits from non-primate mammals. Not only does Sp140L restrict viral infection by preventing the transcription of key viral genes, but also by promoting an ISG response that suppresses cellular pro-proliferative pathways, and curbs the metabolic remodeling required for cellular transformation that is a feature of persistent virus infections.

## Materials and Methods

### EBV Virus Preparation

WT and LPKO viruses were prepared from 293-EBV producer lines and Raji Green Unit titer was obtained as previously described(44). In brief, LPKO virus refers to LPKO^w^ from Szymula et al.(44) which contains a stop codon in exon W2 of every internal W repeat and WT EBV refers to WT^w^ which contains a repaired stop codon in one W repeat found in the parental B95-8 virus that (unlike B95-8) allows robust EBNA-LP expression (94).

### PBMC Isolation and Total/Naïve B and Pan B Cell Enrichment

Adult buffy coats were obtained from the Gulf Coast Regional Blood Center (Pro00006262) and Cord Blood was obtained from the Carolinas Cord Blood Bank (Pro00061264) PBMCs were isolated as previously described (53). Total B cells were isolated using the EasySep Human Pan-B Cell Enrichment kit (STEMCELL Technologies #19554) or MojoSort Human Pan B Isolation kit (Biolegend # 480082). To obtain isolated naïve and memory B cells for SeahorseXF, the EasySep Human Memory B Cell Isolation kit (STEMCELL Technologies #17864) was used. And, to isolate only naïve B cells for the knockout screen and RNA-sequencing, the EasySep Human Naïve B cell isolation kit was used (STEMCELL Technologies, #17254). Purity of isolated cells was assessed by flow cytometry using antibodies to CD19, CD27, and IgD.

### EBV Infection

Isolated B cells were infected by adding WT or LPKO virus at a ratio of 0.2 Raji Green Units per cell and incubating at 37°C for 1 hour. Following incubation, cells were pelleted and resuspended in Roswell Park Memorial Institute (RPMI) medium 1640 supplemented with 20% heat-inactivated fetal bovine serum (FBS) (Corning) (R20 media) at approximately 3 million cells/mL.

### Collection of scRNAseq samples, library preparation, sequencing, and processing

Samples were viably frozen at each time point in 90% FBS + 10% DMSO and stored in liquid N_2_. Cryopreserved samples were then thawed simultaneously and enriched for viable cells by Ficoll gradient. Libraries from 10,000 cells were then prepared in house using the 10x Chromium GEM-X Single Cell 3’ Gene Expression kit (10x Genomics,1000691). Libraries were pooled and sequenced by the Duke Sequencing and Genomic Technologies core facility on the NovaSeq 6000 S2 at 50 base pair read depth, with paired end reads. Reads were mapped to the human genome and type 1 EBV genome as previously described(54).

### scRNAseq QC Filtering and Analysis

scRNAseq analysis was performed in R using package Seurat (95). QC filtering and clustering of samples into subpopulations was performed as previously described(54). To identify the biological features of each cluster, gene ontology analysis was performed using clusterProfiler (96) and the top 100 genes differentially expressed in the previously identified B95-8 subpopulations (54) were compiled and used to add module scores for each subpopulation to the clusters identified in this experiment.

### Flow Cytometry

For analysis of B cell isolation purity (CD19, Biolegend, 302212) and CD23 upregulation (Biolegend, 338516), naïve B cells isolation purity (IgD Biolgened, 348210, CD27 Biolegend 302824), CD46 knockout efficiency (Biolgened, 352405), and ICAM1 expression (Biolegend, 353116) flow cytometry was performed on the BD FACSCanto II (BD Biosciences) and samples were prepared as previously described(53). When quantifying outgrowth of CD46 negative cells, the high throughput sampler was used to collect precise sample volumes.

Flow cytometry was also performed to validate the scRNA-sequencing data using the identified markers using spectral flow cytometry on a Cytek Aurora (Cytek Biosciences). Isolated Pan B cells were stained with CellTraceViolet (ThermoFisher, MPK1096) at 0.1 µM in PBS for 20 minutes at 37°C with gentle mixing every 5 minutes. Staining was then quenched by adding 4x volume of R20 media and incubating for 5 minutes at 37°C. An aliquot of cells was also infected that were not stained with CellTraceViolet to use as a staining control. After pelleting, cells were infected with virus. At each time point collected, cells were first washed in PBS and stained with LiveDead Blue (ThermoFisher, L23105) for 20 minutes in the dark. Samples were then washed with FACS buffer (PBS + 2% FBS) and stained with antibodies as above (CCR6 Invitrogen, 12-1969-42, CD23 Biolegend, 338516, ICAM1 Biolegend 353114, CD38 Biolegend 303529). Unstained cells were used for reference controls for unmixing at each time point, along with single stained compensation beads (BD Biosciences, 552843). To ensure accurate staining and analysis, fluorescence minus one controls, isotype controls, and single stained samples were also prepared for each time point. All flow cytometry data was analyzed in FlowJo (FlowJo, LLC).

### SeahorseXF Assay

Four days post infection, naïve and memory infected B cells were collected. Using flow cytometry, CD19 and CD23 expression was assessed to ensure comparable infectivity between WT and LPKO.

One day prior to the assay, the XFe96 sensor cartridge was submerged in tissue culture grade water overnight at 37°C in a non-CO_2_ incubator, along with XF calibrant (Agilent, 100840-00). The following day, the sensor cartridge was then placed in a utility plate containing the pre-warmed XF Calibrant for 1 hour in the non-CO_2_ incubator before running the assay.

A 96-well Seahorse XF plate was coated with poly-D-lysine to keep suspension cells adhered to the bottom of the wells. Cells were then harvested to the plate at 250,000 cells per well in 50 µL XF Base Medium (RPMI supplemented with 10 mM glucose, 1 mM sodium pyruvate, and 2 mM L-glutamine) (Agilent, 103681-100) and adhered to the plate by briefly centrifugation. 130 µL warm assay media was then added to each well and placed in the non-CO_2_, 37°C incubator for 1 hour.

The XF Real-time APT Rate Assay was used (Agilent, 103592-100). Ports were loaded with oligomycin and Rotenone/Antimycin A at working concentrations of 1.5 µM and 0.5 µM respectively. The calibrated sensor was then placed in the plate and the assay was then run on the Seahorse XF Pro Analyzer (Agilent).

### Bulk RNASeq Sample Collection

RNAseq libraries were prepared from samples 8 days post infection. Pan B cells from three donors total were isolated, however, due to low yield, two of the three donors were pooled together at the time of infection for a total of two Pan B pools infected with separately with WT and LPKO virus. A Ficoll gradient was used to remove dead cells from the harvested samples.

RNA-seq libraries were also prepared 5 days post infection for two donors in which naïve B cells were isolated, transfected with guides targeting CD46, CD46 and SP100, or CD46 and SP140L prior to infection with LPKO or WT virus. Due to low cell counts, cells were not sorted and therefore contain a mixed population of CD46 negative and positive cells (**Fig. S6A**), although the majority of cells were CD46 negative.

### Bulk RNASeq Library Preparation and Analysis

RNA was isolated using the Qiagen RNeasy mini extraction kit including on-column DNase digestion (74104). Libraries were prepared using the NEBNext Ultra II RNA Library Prep kit (E770S) and sequenced as previously described(53). QC of prepared libraries was assessed using a TapeStation (Agilent). Reads were aligned to hg38 (Day 8 bulk RNAseq samples) or hg38 with the Type 1 EBV genome (Day 5 knockout RNAseq samples) using Hisat2. Samtools was then used to generate bam files. DESeq2 was used to generate a ranked list of differentially expressed genes for Gene Set Enrichment Analysis (GSEA) (97).

### Knockout Screen in EBV Infected Naïve B Cells

Following naïve B cell isolation as described above, cells rested overnight in R20 at 37°C. For transfection, Cas9/RNP complexes were prepared from TrueCut Cas9 (ThermoFisher, A36499) and three sgRNAs targeting the gene of interest, and one guide targeting the non-essential cell surface proxy marker CD46 to reduce DNA damage after confirming efficiency of the single CD46 guide (Synthego) (**Table S2**). Cells were washed and resuspended in Buffer T (Neon Transfection System) as previously described (53). 400,000 cells were transfected with Cas9/RNP complexes using the ThermoFisher Neon Transfection System at 2150 V, 20 ms, 1 pulse and resuspended in 200 µL R20 in a 96 well V-bottom plate and incubated at 37°C. One hour after transfection, cells were infected with WT or LPKO virus as described above. Following infection and removal of virus, cells were resuspended in 200 µL R20 and moved to a flat bottom 96 well plate.

Precise sample volumes were used for flow cytometry at each time point to calculate total cell numbers. Two days post infection, cells were stained with anti-CD19 (Biolegend, 302212) to quantify total number of cells per well for normalization. 7-, 14-, 21-, and 28-days post infection, samples were stained with anti-CD46 (Biolegend, 352409) to quantify outgrowth of edited cells. As infected cells expanded, fresh R20 was added in precise volumes to maintain optimal cell density and allow for calculation of total cells per sample. Generation of LCLs was determined by observations including cell clumping and media color change indicative of proliferation and ability to passage or expand cells *in vitro*.

### Trans-complementation Assay

This assay was performed as described previously (53). Following PBMC isolation, cord blood was obtained from the Carolinas Cord Blood Ban (Pro00061264). B cells were isolated by Pan-B cell isolation (STEMCELL Technologies #19554). Isolated B cells were transfected with the trans-complementation plasmid (ori-P based episomal plasmid in backbone pCEP4) prior to infection with LPKO or WT virus. The trans-complementation plasmid encoded tdTomato followed by a P2A cleavage site, and either empty vector, wild type EBNA-LP, or mutant EBNA-LP in which the Y domain leucines (Leu276, Leu780, and Leu781), of the sequence LXXXLL were modified to alanine by site-directed mutagenesis (Agilent, 200521). Outgrowth of tdTomato positive cells was assess by flow cytometry each week until 35 days post-infection, with compensation for the viral-encoded GFP. At this final timepoint, wells were considered transformed into tdTomato positive LCLs if cells were both tdTomato-expressing and contained proliferating cells passaged in culture.

### Prediction of Protein Structure by AlphaFold3

Protein structures were predicting using the AlphaFold server (https://alphafoldserver.com) using default settings. To predict the structure of a single molecular of EBNA-LP, the sequence of wild type EBNA-LP (UniProt Q8AZK7) encoding 4 W domains was used for input, and the representative model 0 was selected for visualization using PyMOL (98). For EBNA-LP/Sp140L (UniProt Q9H930) and EBNA-LP/Sp100 (UniProt P23497) complexes, jobs were submitted with 2 molecules of each protein. Model 0 for each complex was representative of predicted models and selected for visualization. For complexes containing SP140L, isoform 2, which excludes the frequently skipped exon 2(29), was used. For Sp100, the Sp100C isoform was used as it contains the most similar domain architecture to Sp140L.

### Co-Immunoprecipitation in 293Ts

293T cells transfected with 20 ug total DNA, using pSG5 empty vector as filler DNA. Constructs expressing EBNA-LP encode synthesized cDNA (GENEWIZ) encoding four copies of the identical W domain with optimized codon usage to avoid recombination and were cloned by gateway cloning (ThermoFisher 11789020, and 11791020) into the vector pSG5-FLAG-Gateway (a gift from Eric Johannsen) to include an N-terminal FLAG tag. DNA encoding Sp100A was from plasmid YFP-Sp100A(K297A)-C1 (Addgene plasmid # 134552). The sequence encoding Sp100A was inserted into the vector pCDNA3.1-HA, adding an N-terminal epitope tag, by restriction digest cloning. Site-directed mutagenesis cloning was used to restore Sp100A sequence to wild type (A297K).

FLAG immunoprecipitations were performed as described previously (53) with 4 ug Anti-FLAG M2 antibody (Sigma-Aldrich, F1804). Beads were eluted in 1X LDS sample buffer (GenScript, M00676-250) in lysis buffer at 37°C for 5 min followed by rotation at room temperature for 5 min. Immunoprecipitation blots were imaged using primary antibodies EBNA-LP JF186 (purified in house, used at a final dilution of 10 ug/uL) and Sp100 polyclonal antibody (Thermo PA5-78177, 1:750). Veriblot for IP Detection Reagent (HRP, abcam ab131366) was used for secondary antibody at 1:1000 dilution in 5% milk in TBST. Sp100 input western blot was imaged using SuperSignal™ West Pico Chemiluminescent Substrate (Thermo Scientific 34080), while all other western blots were imaged using SuperSignal™ West Femto Maximum Sensitivity Substrate (Thermo Scientific 34096).

### Generation of Recombinant HVS Virus

A BAC clone of the HVS A11-Sac strain (which lacks the HVS transforming genes) (99) was modified using RecA-mediated recombineering to replace the AgeI-BlpI fragment of dsRed1 ORF with a mNeonGreen-T2A-Luciferase fusion. This generated two BAC clones (A and B) that were each recombineered to delete the ORF3 open reading frame, introducing a STOP codon (alongside SalI and Cla I restriction sites) after the 5^th^ amino acid of ORF3, and deleting all but the last 237 nucleotides of the ORF3 coding sequence, thereby generating two independent HVS-ORF3KO BACs (A-ORF3 KO and B-ORF3 KO) (as validated in **Fig. S8A**).

### HVS Virus Preparation

Herpesvirus saimiri BAC is transfected into – using a peptide-lipofectin complex (100) – or inoculated onto Owl Monkey kidney (OMK) cells. Supernatant is harvested when the cell monolayer is largely destroyed (typically after 4-7 days) and centrifuged to remove cell debris.

### HVS Genome Quantification

Virus stock was lysed by lysis buffer after DNase treatment. Then virus lysate was subjected to quantify viral DNA by using Kapa SYBR Fast Universal qPCR kit (KK4602, SLS) with a pair of hygromycin gene primer.

### Generation of *CD46*, *SP100*, and *SP140L* Knockout Fibroblast Cell Lines

Human Foreskin Fibroblasts (HFF) and human lung fibroblast cell line MRC-5 (MRC5) were maintained in Dulbecco’s modified Eagle’s medium (DMEM) (Gibco, Grand Island, NY, USA) supplemented with 10% heat-inactivated fetal bovine serum (Gibco, Grand Island, NY, USA) at 37°C in a 5% CO2. For transfection, Cas9/RNP complexes were prepared in buffer T whereas cells were washed and resuspended in buffer R (Neon transfection system). 2x10^5^ cells in 5 ul buffer R were mixed with Cas9/RNP complexes and transfected with the neon transfection system at 1650 V 10 ms, 3 pulses. Then the transfected cells were resuspended in 300 µl of 15% DMEM in 12 well plate and incubated at 37°C. When cell confluency reached 80-90%, a quarter of the well was stained with anti-CD46 antibody (Biolegend, 352401) to measure CD46-knockout efficiency by flow cytometry as described above.

### Immunofluorescence in Fibroblast Knockout Lines

HFF cells were seeded onto coverslips. At 24 hours post HVS infection, the coverslips were fixed with 4% PFA for 15 minutes and permeabilized with 0.5% Triton X-100 in PBS for 15 minutes. Coverslips were blocked with 10% FBS in PBS for 1 hour. The coverslips were incubated with the anti-Sp100 (Invitrogen, PA5-78177) for 1 hour followed by Alexa Fluor 546 conjugated anti-rabbit IgG (Invitrogen, A11071) for 1 hour. The coverslips were mounted with ProLong® Gold Antifade Reagent with DAPI (Cell Signalling Technology, 8961). Images were obtained using the EVOS cell imaging system.

### Knockout Screen in HVS Infected Fibroblast Cell Lines

Fibroblast cells were seeded 100 ul of 3x10^5^ cells/ml triplicate wells of a white 96 well clear-bottom plate. HVS WT (A and B) and HVS ORF3 KO (A-ORF3 and B-ORF3) were used to infect knockout cell line at MOI of 1000 genomes/ml. After 24 hours, luciferase activity was measured as a proxy for HVS transcription: cell media was removed and replaced with 50 µl fresh media, to which 50 µl Steady-Glo® Luciferase Assay System reagent (Promega, E2510) was added, and after 5 minutes, luminescence was measured on a FluoStar OMEGA plate reader (BMG Labtech).

### Immunofluorescence in EBNA-LP trans-complemented LCLs

Samples were pelleted and resuspended in 200uL of 1X PBS per slide. Cells were spun onto slides using a Cytospin 4 (1000rpm for 8 min). Cells were fixed with 4% paraformaldehyde in PBS for 15 min at 4°C. Cells were then washed three times in 1X PBS, permeabilized in 0.5% Triton X-100 in 1X PBS for 15 min at RT, and blocked with 5% normal goat serum in 0.2% Triton X-100 in PBS in a humid chamber for 1 hour at RT. Primary antibodies were then added to blocking buffer (JF186 purified from hybridoma supernatant in-house, used at a final dilution of 10 µg/μL) and incubated on slides with parafilm in humid chamber overnight at 4°C. Slides were washed four times in 1X PBS, incubated with secondary antibody in blocking buffer (AlexaFluor 488 goat anti-mouse IgG 1:250, Cat. No A11001) for 2 hours at RT and washed 4 times in 1X PBS. Slides were mounted with Vectashield Vibrance Antifade mounting medium with DAPI (Vector Laboratories H-1800) and imaged as Z stacks on the Olympus IX81 microscope using 60X oil objective. Images were processed with Fiji (ImageJ).

### Knockout Validation

Knockout cell lines were validated by genomic DNA (gDNA) sequencing at targeted cut sites. For LPKO LCLs, the total population of LCLs (regardless of CD46 expression) was used to collect samples for gDNA – as if targets have the ability to rescue LPKO infection, we expect that all surviving cells are edited. Cells were pelleted and washed in PBS. gDNA was isolated (K182002, ThermoFisher) and primers spanning cut sites (**Table S2**) were used to amplify region of interest. gDNA from control (CD46 only) conditions were used for comparison. PCR product was sequenced by Sanger Sequencing. The chromatogram files generated from sequencing were used as input in the Synthego Inference of Crisper Edits Analysis tool (https://www.synthego.com/products/bioinformatics/analysis), with CD46 only control cells used for comparison. This was used to generate a knockout score, indicative of the percentage of sequences, a direct reflection of the percentage of cells, that have indels that are highly likely to result in lost expression of the targeted gene by non, either from frameshifting indels or indels larger than 21 base pairs that would cause either non-sense mediated decay or loss of function mutations.

### Western Blotting

Western blots were performed as previously described (53).

### Resource Availability

All sequencing data is publicly available from the NCBI’s Gene Expression Omnibus (GEO) under accession numbers: GSE282376, GSE282377, and GSE282400.

## Acknowledgements

This work was supported by NIH Grant R01CA140337 (to M.A.L), MRC grant MR/L008432/1 (to R.E.W.), and F31DE031509 (to J.M.C.). We acknowledge Ashely P. Barry for reagent generation and technical assistance with the SeahorseXF assay, Elliott SoRelle for assistance with scRNAseq analysis, Gillian Horn for assistance with design of flow cytometry panels, Neha Shaw for technical assistance, Michael di Franco for pilot HVS experiments, Johanna Veldman for historic Sp140L experiments, and Nicolás Reinoso-Vizcaino for helpful discussions. We wish to thank the Duke University School of Medicine Sequencing and Genomic Technologies Shared Resource for sequencing services and the Duke Cancer Institute Flow Cytometry Facility for instrumentation.

