## Supplemental Figures for "Sp140L Is a Novel Herpesvirus Restriction Factor"

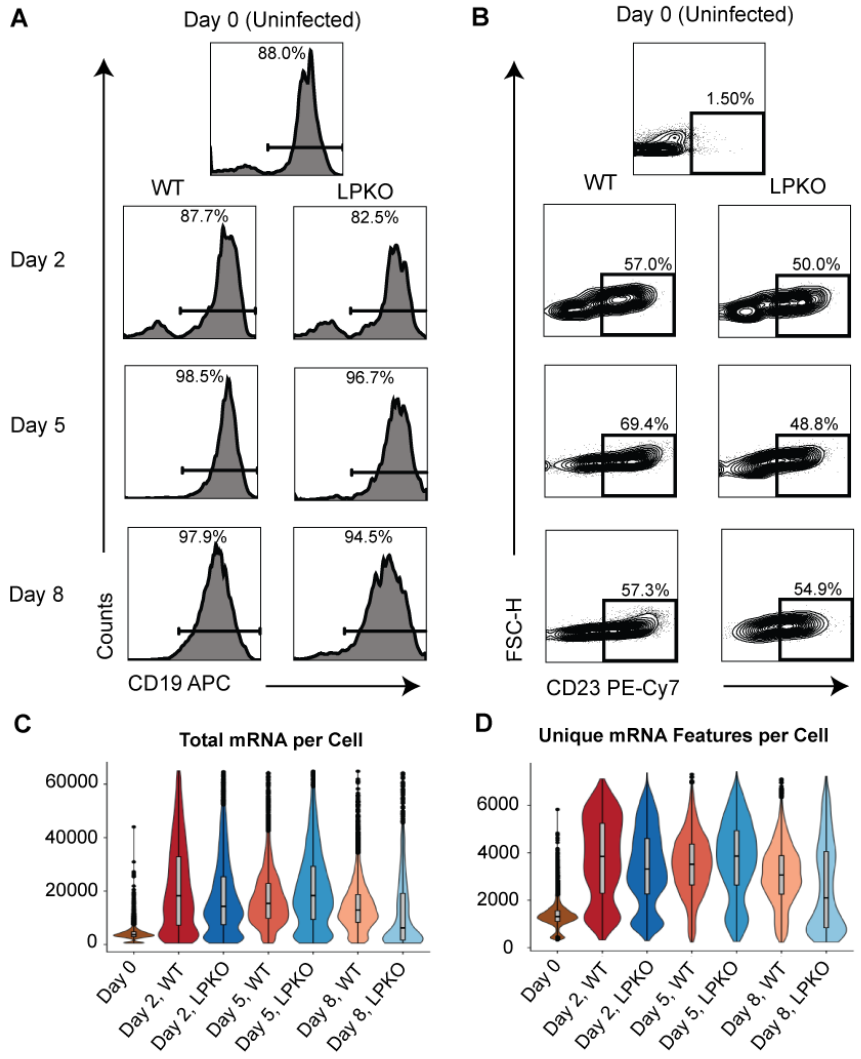


**Fig. S1. Quality control of samples and scRNAseq libraries. A.** Histogram of CD19 expression for each sample prior to collection. **B.** CD23 expression for each sample prior to collection. **C.** Total mRNA molecules read per cells in each sample. **D.** Total unique mRNA molecules per cell in each sample.

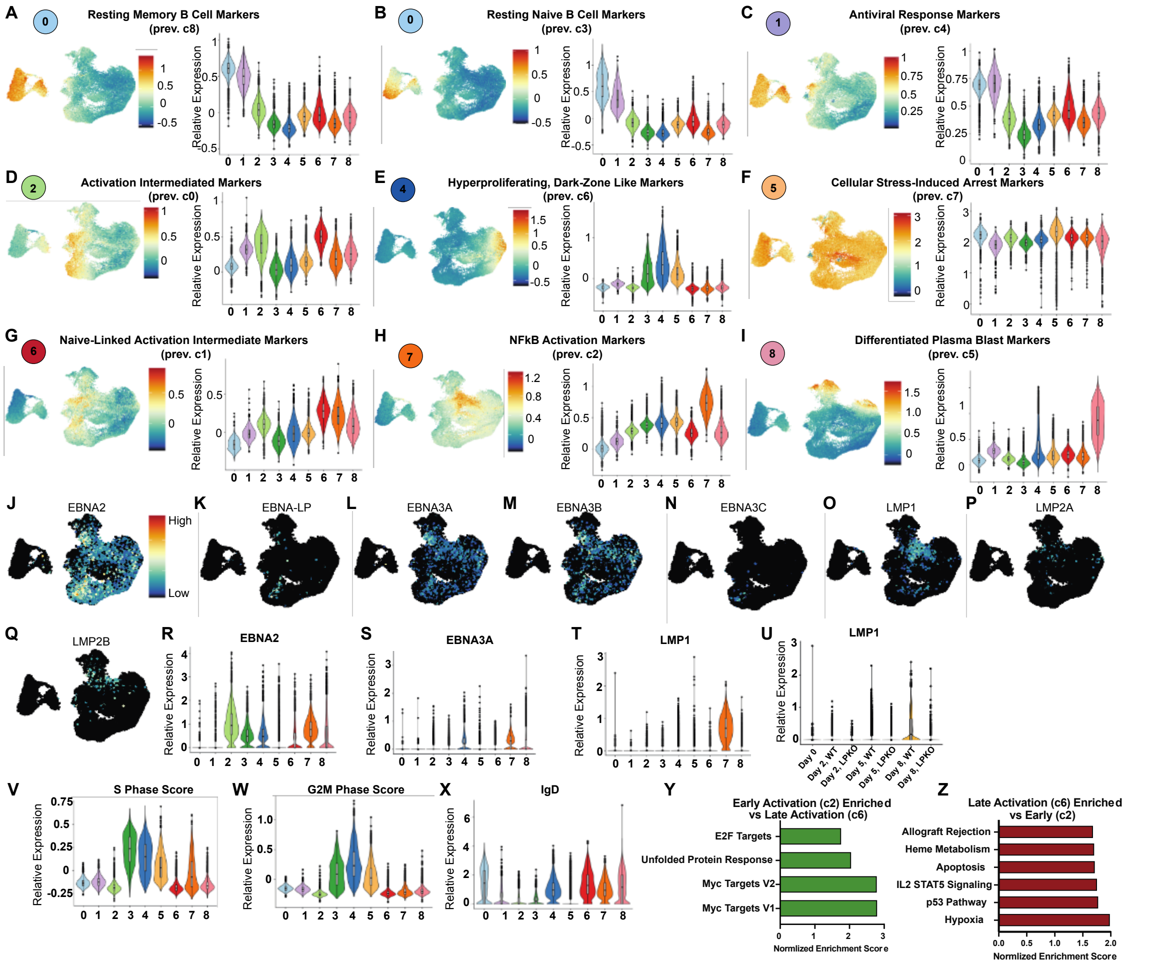


**Fig. S2. Identified clusters correlate with EBV-induced cell states, viral gene expression, and cell cycle phase.** Module scores based on top 100 differentially expressed genes for each previously identified B95-8 associated subpopulation**.** Umaps highlight cells with highest expression of indicated markers. Violin plot indicates relative expression of markers across clusters. The corresponding cluster number in prior publications is indicated for reference (2). **A.** Resting Memory B cell Markers. **B.** Resting Naïve B cell Markers. **C.** Sensing/Antiviral Response Markers. **D.** Activation Intermediate Markers. **E.** Hyperproliferation/Dark Zone Like Markers. **F.** Cellular Stress-Induced Arrest Markers. **G.** Naïve-linked Activation intermediate Markers. **H.** NFκB-High/Light Zone Markers. **I.** Differentiated/Plasmablast State Markers. **J**. Umap of viral latency gene EBNA2 across all cells. **K.** EBNA-LP. **L.** EBNA3A. **M.** EBNA3B. **N.** EBNA3C. **O.** LMP1. **P.** LMP2A. **Q.** LMP2B. **R.** Violin plot of relative expression of viral gene EBNA2 across cluster. **S.** EBNA3A. **T.** LMP1. **U.** Violin plot of LMP1 expression by sample. **V.** Relative enrichment of S phase score across clusters. **W.** G2M phase score. **X.** Relative enrichment of IgD across clusters. **Y.** Significantly enriched (FDR q-value < 0.05) GSEA Hallmark pathways in Early Activation state compared to Late. **Z.** Enriched in Late compared to Early.

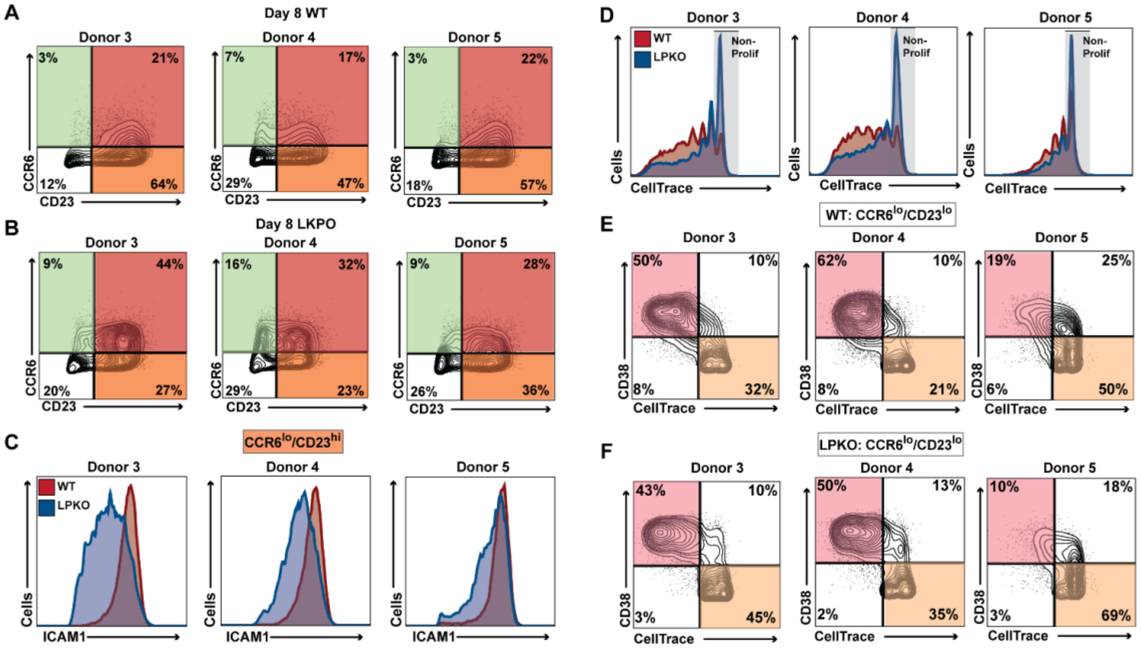


**Fig. S3. Validation of scRNAseq data by flow cytometry in additional donors.** Flow staining for CCR6 and CD23 8 days post-infection in additional donors infected with WT (**A**) or LPKO virus (**B**). Quadrant color corresponds to cell states that can be distinguished as depicted Figure 2E. **C**. Histograms comparing expression of ICAM1 protein (proxy for LMP1) between WT and LPKO infected cells in CCR6$^{lo}$/CD23$^{hi}$ population. **D.** Histogram of CellTrace stain, which is diluted during each cell division. Grey area indicates cells that have not undergone division. Separation of CCR6$^{lo}$/CD23$^{lo}$ populations in WT (**E**) and LPKO (**F**) infection by CD38 and CellTrace to distinguish Differentiated cells from Stressed cells. Histograms are scaled as percent of maximum count (modal).

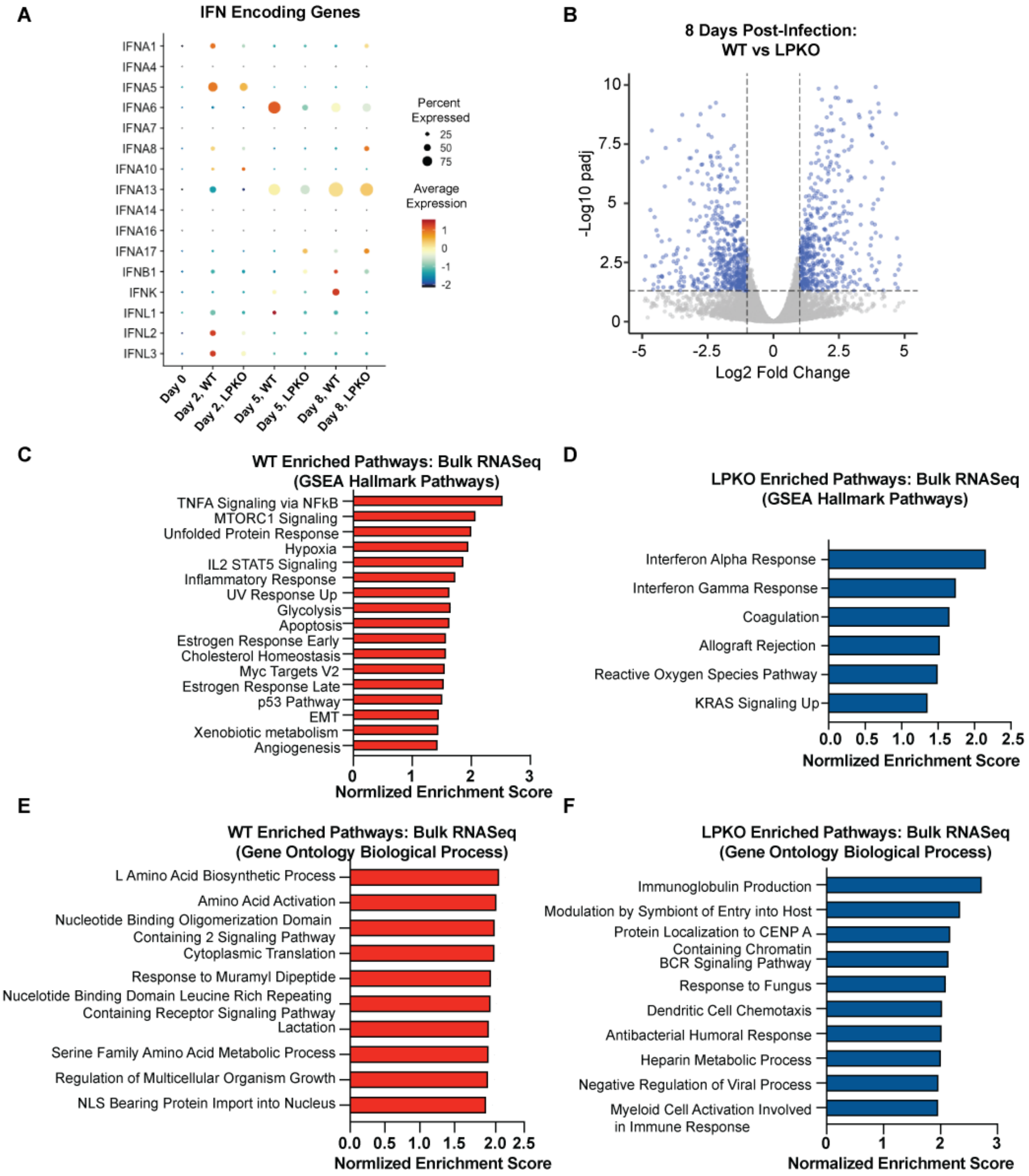


**Fig. S4. Differential gene expression analysis in bulk RNAseq data 8 days post infection corroborates scRNA-seq data. A.** Expression of genes encoding IFN detectable in scRNAseq time course samples. **B.** Volcano plot of differentially expressed genes between WT and LPKO samples. Gene sets enriched in WT infected cells compared to LPKO (**C**) and LPKO infected cells compared to WT (**D**) by GSEA analysis using Hallmark pathways by bulk RNA-seq. All gene sets with false discover rate (FDR q-value) less than 0.05 are plotted. Higher absolute value of Normalized Enrichment Score correlates with stronger enrichment. Gene sets enriched in WT infected cells compared to LPKO (**E**) and LPKO compared to WT (**F**) by GSEA analysis using Gene Ontology Biological Processes. The top 10 non-redundant enriched pathways with the highest Normalized Enrichment Score and FDR q-value less than 0.05 are shown.

**A**

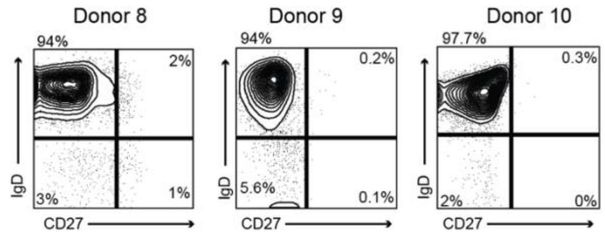


**B**

Day 28 LPKO LCLs

| Donor 8 | | | Donor 9 | | | Donor 10 | | |
|---|---|---|---|---|---|---|---|---|
| Target | Well | KO Score | Target | Well | KO Score | Target | Well | KO Score |
| SP100 | A | 100 | SP100 | A | 100 | SP100 | A | 100 |
| | B | 100 | | B | 100 | | B | 100 |
| | C | 98 | | C | 100 | | C | 100 |
| SP140L | A | 100 | | D | 100 | | D | 100 |
| | B | 99 | SP140L | A | 99 | SP140L | A | 100 |
| | C | 98 | | B | 100 | | B | 100 |
| | D | 0 | | C | 100 | | C | 100 |
| | | | | D | 88 | | D | 100 |
| | | | DAXX | A | 100 | SP140 | A | 100 |
| | | | | B | 100 | | | |

**C**

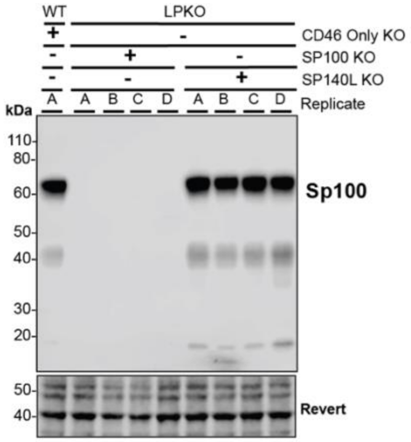


**D**

7 Week LPKO LCLs

| Donor 9 | | | Donor 10 | | |
|---|---|---|---|---|---|
| Target | Well | KO Score | Target | Well | KO Score |
| DAXX | C | 0 | CD46 | A | NA |
| | D | 86 | DAXX | A | 0 |
| SP110 | A | 0 | | B | 100 |
| | B | 0 | | C | 100 |
| | C | 0 | | D | 84 |
| | | | SP110 | A | 0 |
| | | | | B | 2 |
| | | | SP140 | B | 90 |
| | | | | C | 100 |
| | | | PML | A | 67 |
| | | | | B | 4 |
| | | | | C | 80 |
| | | | IFI16 | A | 96 |

**E**

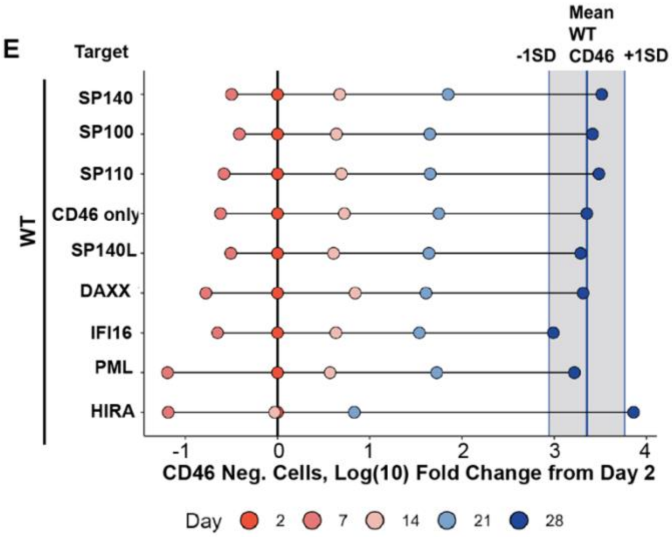


**F**

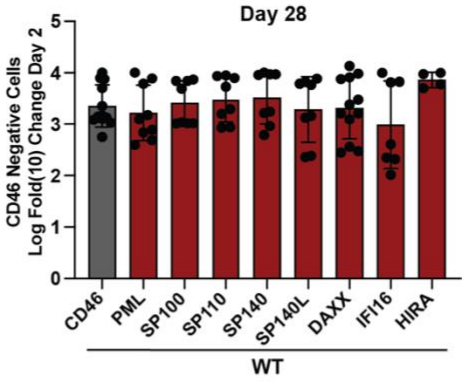


**Fig. S5. Loss of *SP100* and *SP140L* does not impact outgrowth of WT EBV infected naïve B cells. A.** Purity of isolated CD19 positive naïve B cell fractions for donors in screen. **B.** Knockout score for each LPKO LCL generated by 28 days post infection, indicative of the percentage of cells with the target gene knocked out based on indel frequency. **C.** Log(10) Fold Change from Day 2 of the total number of CD46 negative cells for each condition 2, 7-, 14-, 21-, and 28-days post-infection. Dark turquoise line represents the mean number of CD46 negative cells in CD46 only control WT infected cells 28 days post infection. **D.** Total number of CD46 negative cells 28 days post infection for each condition plotted as log fold change from 2 days post infection. P values calculated by one-way ANOVA with multiple comparisons. ** indicates p-values <0.01. **E.** Knockout score of LPKO LCLs generated after a total of seven weeks in culture, significantly delayed outgrowth compared to rescued LPKO LCLs and WT LCLs. **F.** Western blot for Sp100 protein expression in LPKO LCLs with *SP100* KO or *SP140L* KO from donor 3. Molecular weight in kDa is indicated.

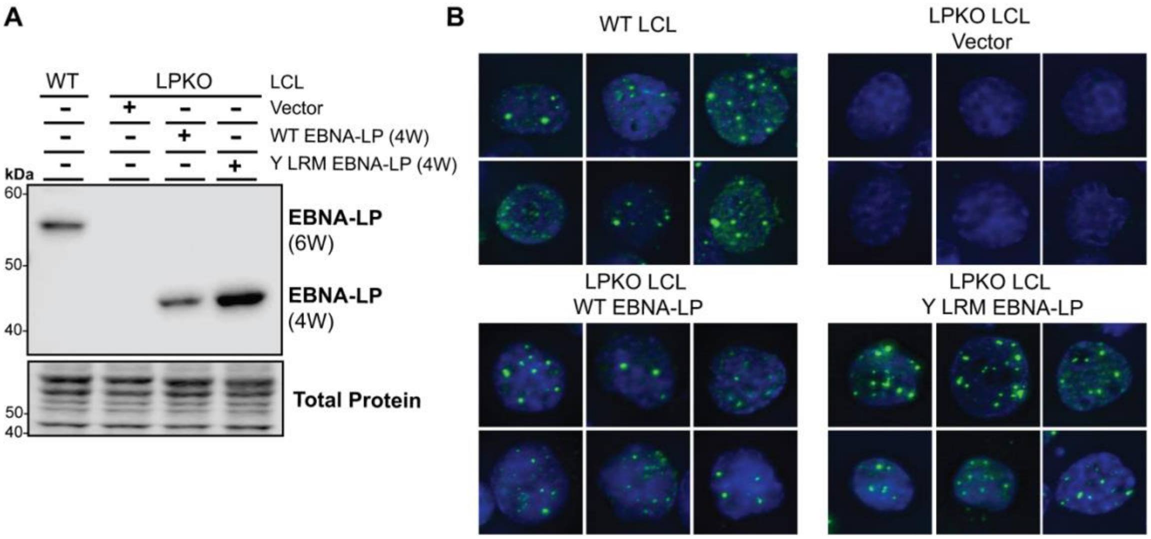


**Fig S6. Y LRM Mutant EBNA-LP displays similar expression and localization patterns as WT EBNA-LP. A.** Western blot analysis of LPKO-infected total B cells trans-complemented with empty vector, wild type, or Y LRM mutant EBNA-LP. LCLs derived from WT virus (WT LCLs) used as control. DNA constructs used for trans-complementation encode 4 repeated W domains (4W) while the virus encodes 6 (6W). **B.** Immunofluorescence of WT and trans-complemented LPKO-infected total B cells. Green indicates staining for EBNA-LP. Blue indicates DAPI staining for nuclei.

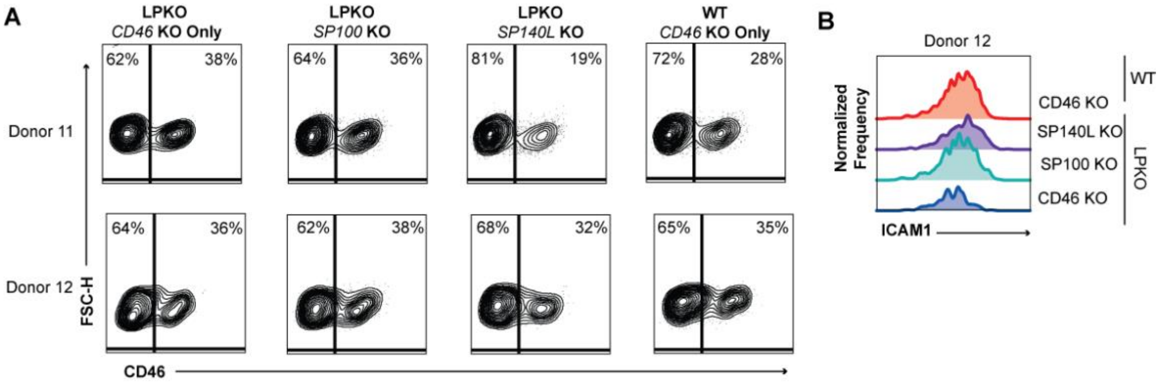


**Fig S7. RNAseq samples for *SP100* and *SP140L* knockout. A.** CD46 negative and positive populations for each RNAseq sample at time of collection. **B.** ICAM1 expression in each sample from second donor not shown in Fig 6.

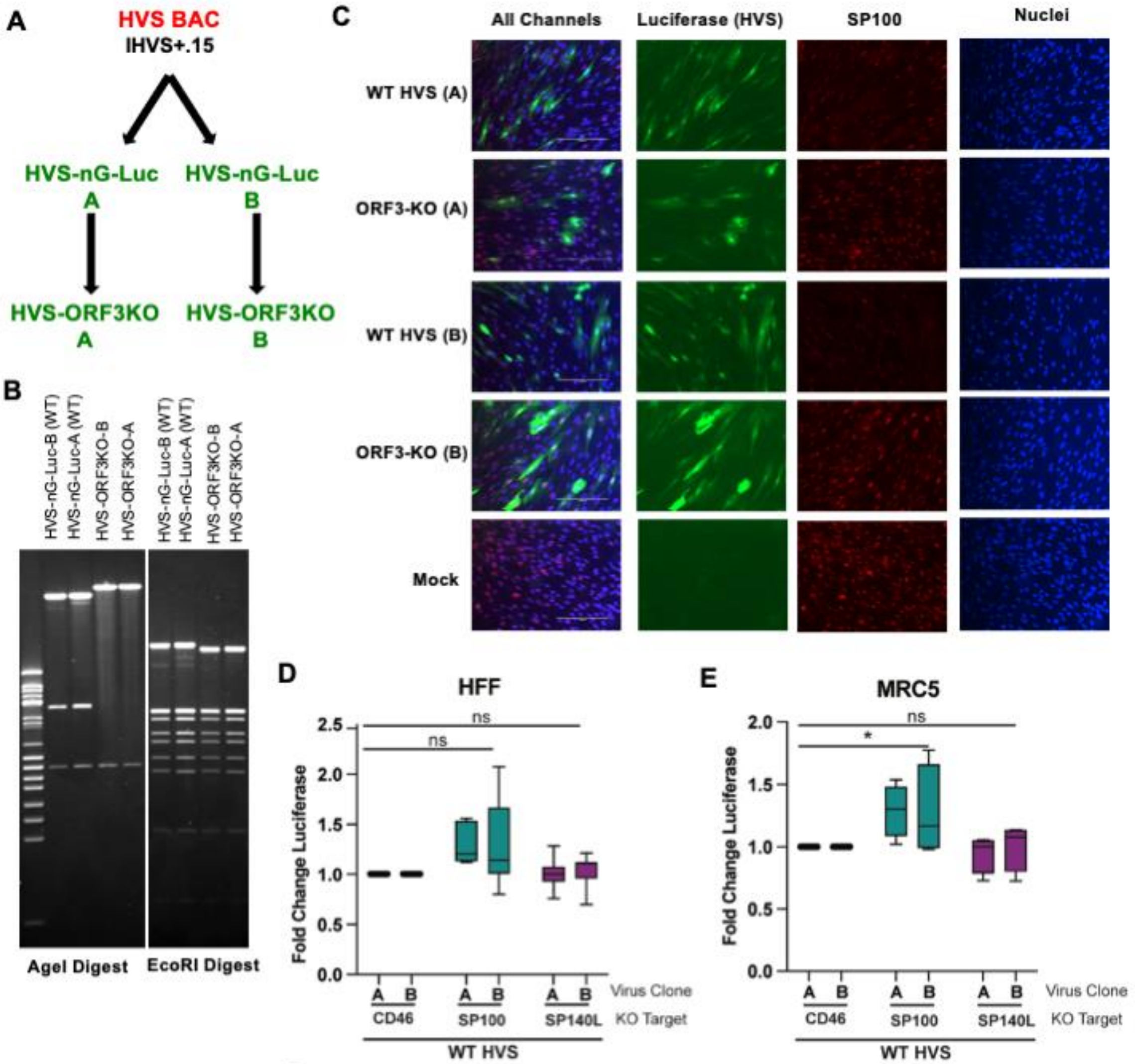


F

| | HFF | MRC5 |
|---|---|---|
| Target | Knockout Score | Knockout Score |
| Sp100 | 95 | 100 |
| Sp140L | 100 | 97 |

**Fig S8. *SP100* and *SP140L* loss has minimal effect on wild type HVS. A.** Flow chart showing recombineering steps to generate the recombinant HVSs used in this study. Green indicates viruses with nGreenLuciferase (nGL-Luc) inserted. **B.** Restriction digest of WT HVS and ORF3-KO BACs confirming insertion of stop codon in ORF3-KO virus genome. **C**. Immunofluorescence of HFF cells infected with WT and ORF3-KO viruses. Green indicates nGreenLuciferase expression encoded in viral genome. Red indicates staining for Sp100. Blue indicates stained DNA. **D.** Relative luciferase expression in HFF knockout cells infected with WT HVS. **E.** MRC5 cells. P values calculated by two-way ANOVA with multiple comparisons. * Indicates p-values <0.05. **F.** Validation of SP100 and SP140L knockout by gDNA sequencing. Knockout score indicates percentage of sequences in a population in which indels likely to result in loss of protein expression were identified.
